# Modulation of xanthophyll cycle impacts biomass productivity in the marine microalga *Nannochloropsis*

**DOI:** 10.1101/2022.08.16.504082

**Authors:** Giorgio Perin, Alessandra Bellan, Dagmar Lyska, Krishna K. Niyogi, Tomas Morosinotto

**Author notes:** Corresponding author: Tomas Morosinotto, Department of Biology, University of Padova, Via Ugo Bassi 58/B, 35131, Padova, Italy., **Email:**. Equal contribution (Giorgio Perin, Alessandra Bellan). **Author Contributions:** TM, conception and design. AB, data collection. GP, collection and critical revision of data. DL and KKN, critical revision of data, manuscript and generation of the *N. oceanica* mutants. GP and TM, writing of the manuscript. All authors, final revision of the manuscript.

## Abstract

Life on earth depends on photosynthetic primary producers that exploit sunlight to fix CO_2_ into biomass. Approximately half of global primary production is associated with microalgae living in aquatic environments. Microalgae also represent a promising source of biomass to complement crop cultivation, and they could contribute to the development of a more sustainable bioeconomy. Photosynthetic organisms evolved multiple mechanisms involved in the regulation of photosynthesis to respond to highly variable environmental conditions. While essential to avoid photodamage, regulation of photosynthesis results in dissipation of absorbed light energy, generating a complex trade-off between protection from stress and light-use efficiency. This work investigates the impact of the xanthophyll cycle, the light-induced reversible conversion of violaxanthin into zeaxanthin, on the protection from excess light and on biomass productivity in the marine microalgae of the genus *Nannochloropsis*. Zeaxanthin is shown to have an essential role in protection from excess light, contributing to the induction of Non-Photochemical Quenching and scavenging of reactive oxygen species. On the other hand, the overexpression of Zeaxanthin Epoxidase, enables a faster re-conversion of zeaxanthin to violaxanthin that is shown to be advantageous for biomass productivity in dense cultures in photobioreactors. These results demonstrate that zeaxanthin accumulation is critical to respond to strong illumination, but it may lead to unnecessary energy losses in light-limiting conditions, and accelerating its re-conversion to violaxanthin provides an advantage for biomass productivity in microalgae.

**Significance Statement:** This work investigates the impact of the xanthophyll cycle in marine microalgae on the trade-off between photoprotection and light-use efficiency. Our results demonstrate that whilst zeaxanthin is essential for photoprotection upon exposure to strong illumination, it leads to unnecessary energy losses in light-limiting conditions and thus accelerating its re-conversion to violaxanthin provides an advantage for biomass productivity in microalgae.

## Introduction

Photosynthetic organisms are the main primary producers on our planet, supporting the metabolism of most life forms, thanks to their ability to exploit sunlight to drive the fixation of CO_2_ into biomass. Approximately half of global primary production is associated with aquatic environments and depends on microalgae, making these organisms essential to sustain life in natural ecosystems (1). Investigating the regulation of photosynthesis is essential both to understand the dynamics of primary productivity in natural ecosystems as well as to pave the way to improve light-to-biomass conversion efficiency and increase crop productivity to respond to an ever-increasing demand for food (2).

In the natural environment, light absorbed by photosynthetic pigments, such as Chlorophyll (Chl), can easily become excessive with respect to the metabolic capacity of the cell, driving the over-reduction of the photosynthetic electron transport chain and consequently the generation of toxic reactive oxygen species (ROS). Photosynthetic organisms evolved several mechanisms regulating light-use efficiency and photosynthetic electron transport to reduce the probability of over-reduction and cell damage (3, 4). Among these mechanisms, Non-Photochemical Quenching (NPQ) drives the dissipation of excited states of Chl (i.e. Chl singlets) as heat, thus reducing the probability of generating ROS. In eukaryotes, NPQ depends both on the generation of a ΔpH across the thylakoid membrane and the presence of specific molecular activators, namely PsbS and/or LHCSR/LHCX, depending on the species (5, 6).

In most eukaryotic organisms, a second major regulatory mechanism of photosynthesis is the xanthophyll cycle. Upon exposure to excess irradiation, the decrease in pH of the thylakoid lumen induces the activation of Violaxanthin De-Epoxidase (VDE) that catalyses the conversion of violaxanthin into zeaxanthin (7, 8). Zeaxanthin contributes to photoprotection both by enhancing NPQ and directly scavenging Chl triplets and ROS (9). In limiting light conditions, zeaxanthin is converted back to violaxanthin by Zeaxanthin Epoxidase (ZEP). The two reactions of the cycle have different kinetics and, while zeaxanthin accumulates in a few minutes after exposure to strong illumination, it takes tens of minutes for ZEP to convert it back to violaxanthin. This slower rate of re-conversion has been suggested to provide more effective photoprotection in nature in case of repeated peaks of excess irradiation due to rapidly changing weather conditions (10).

NPQ and the xanthophyll cycle are important to protect the photosynthetic apparatus from excess irradiation, and they have been shown to contribute to the fitness of photosynthetic organisms in dynamic natural conditions (11). On the other hand, their activity results in the dissipation of a fraction of absorbed energy (12), reducing light-to-biomass conversion efficiency. If constitutively active, thus, they can negatively impact biomass productivity in light-limiting conditions (13). The energy losses due to photosynthesis regulatory mechanisms can be particularly impactful in the case of light fluctuations, when NPQ and the xanthophyll cycle are activated during light peaks and remain active when the illumination decreases. In plants it has been shown that accelerating the kinetics of the xanthophyll cycle can lead to a remarkable increase in photosynthetic productivity in the field (14, 15).

Unicellular algae, like all other photosynthetic organisms, are exposed to light fluctuations in nature and have multiple mechanisms to modulate their photosynthetic efficiency, including NPQ and the xanthophyll cycle (16, 17). Light dynamics are also highly impactful when microalgae are cultivated in photobioreactors for commercial applications, where culture optical density and its mixing generate additional light fluctuations, beyond the natural dynamics (18). In this work, we investigated the impact of the xanthophyll cycle in the heterokont marine microalgae *Nannochloropsis gaditana* and *N. oceanica*, showing the essential role of zeaxanthin in photoprotection from light stress but also demonstrating that a faster re-conversion of zeaxanthin to violaxanthin improves biomass productivity in a light-limited environment, typical of dense cultures of industrial systems.

## Results

### Dynamics of xanthophyll composition in Nannochloropsis

*Nannochloropsis gaditana* cultures, exposed to different light intensities, showed accumulation of antheraxanthin and zeaxanthin following the increase in irradiance, with a corresponding reduction in the content of violaxanthin (Figure 1a). It is worth noting that even when grown in limiting light conditions (i.e. < 150 μmol photons m^−2^ s^−1^, (19)], *Nannochloropsis* cells showed a small but detectable presence of zeaxanthin (>2%, Figure 1a), different from plants or other eukaryotic microalgae, where zeaxanthin is normally not detectable in low light (20). Cells exposed to high light (1000 μmol photons m^−2^ s^−1^) for different time intervals showed a progressive increase in antheraxanthin and zeaxanthin with a corresponding decrease in violaxanthin (Figure 1b and supplementary Table S1). Vaucheriaxanthin and β-carotene, the other major carotenoids detected, instead did not change in response to the treatment with excess light (Supplementary Table S1), all results fully consistent with the activation of the xanthophyll cycle induced by the strong illumination. Antheraxanthin content reached a maximum after 15 min, while zeaxanthin accumulation continued to increase, not reaching a saturation even after 2 h of high light treatment (Figure 1b).

**Figure 1.**
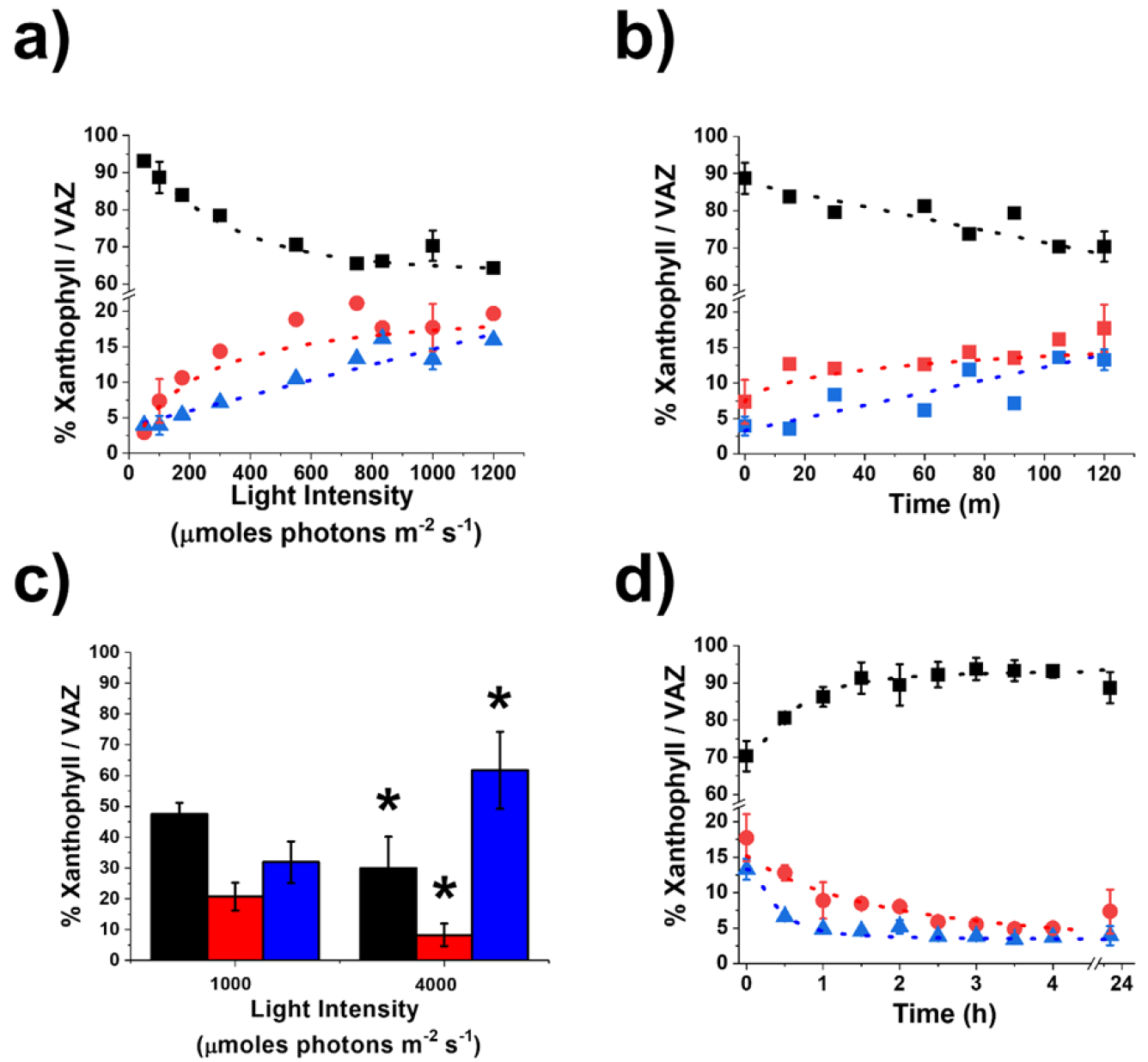
Dynamic of xanthophyll cycle in *Nannochloropsis gaditana*. Light (a) and time (b) dependence of xanthophylls accumulation in *Nannochloropsis gaditana* previously cultivated at 100 μmol photons m^−2^ s^−1^. c) The same Nannochloropsis gaditana cells were also exposed to extreme illumination (1000 and 4000 μmol photons m^−2^ s^−1^) for 2 h, removing CO_2_ to maximize photosynthesis saturation. d) Time-dependent relaxation of xanthophylls after exposure at 1000 μmol photons m^−2^ s^−1^ for 2 h, as in panel b. Data are fitted with logistic functions and are expressed as percentage of each xanthophyll molecule over their sum (violaxanthin, antheraxanthin and zeaxanthin, VAZ). Black, violaxanthin; red, antheraxanthin; blue, zeaxanthin. Asterisks in panel c indicate statistically significant differences in the xanthophylls content between 4000 and 1000 μmol photons m^−2^ s^−1^ (One-way ANOVA, p-value<0.05). Data are expressed as average ± SD of three independent biological replicates.

Cells were also treated with extreme, non-physiological, light intensity (4000 μmol photons m^−2^ s^−1^ while also removing CO_2_ supply, Figure 1c), to maximize light excess. This resulted in a further accumulation of zeaxanthin that reached in the most extreme case 60% of the VAZ pool (Figure 1c), showing that this organism has a very large reservoir of violaxanthin convertible to zeaxanthin. To investigate xanthophyll cycle relaxation dynamics, *Nannochloropsis gaditana* cells treated with 1000 μmol photons m^−2^ s^−1^ for 2 h to induce zeaxanthin biosynthesis, were afterwards exposed to dim light (Figure 1d). Dim light was preferred to dark because the former is expected to increase the amount of photosynthesis products, such as O2 and NADPH, that are required by the epoxidation reaction catalyzed by ZEP (21). Zeaxanthin and antheraxanthin synthesized during the high light treatment were fully re-converted to violaxanthin after approximately 4 h (Figure 1d).

### Impact of xanthophyll dynamics on Non-Photochemical Quenching

Exposure to saturating illumination also activates a photoprotection mechanism, called NPQ, that can be quantified by monitoring chlorophyll fluorescence *in vivo* (see Materials and Methods for details). In *Nannochloropsis*, NPQ activation reaches saturation after approx. 10 min of exposure to saturating illumination (Figure 2a). In *Nannochloropsis*, NPQ is strongly influenced by zeaxanthin synthesis, as shown by treatment with a VDE inhibitor (i.e. DTT) that causes a strong reduction of its activation (Supplementary figure S1).

**Figure 2.**
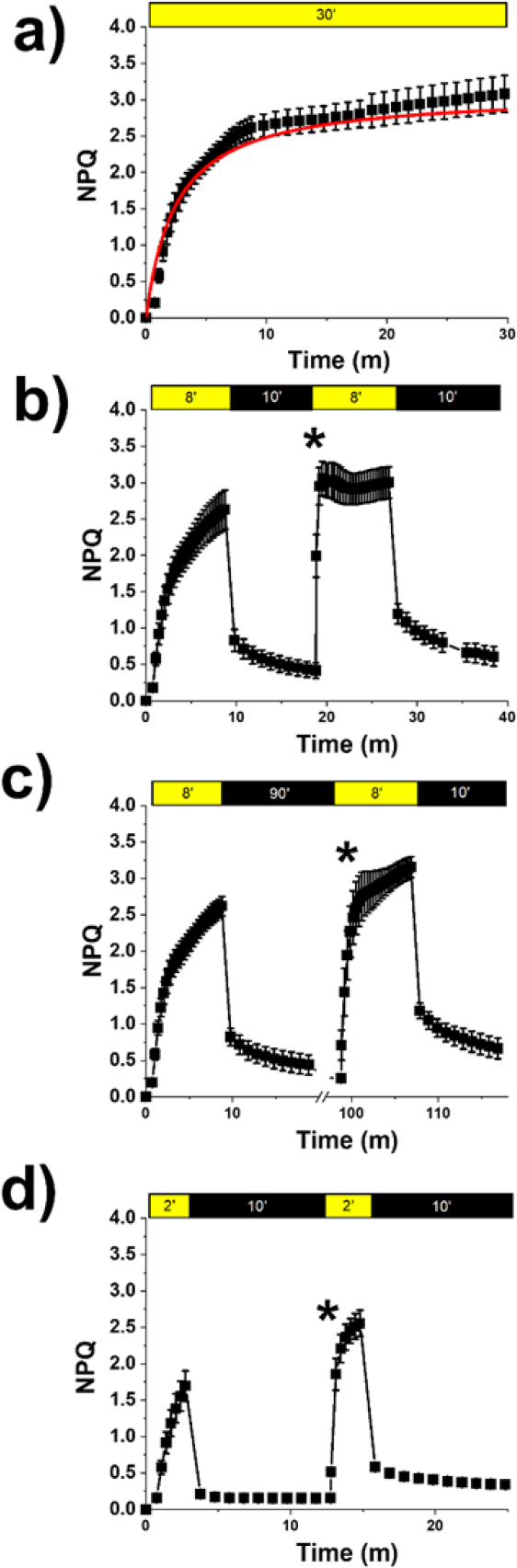
Influence of zeaxanthin on Non-Photochemical Quenching. NPQ kinetics calculated from chlorophyll fluorescence upon exposure of *Nannochloropsis gaditana* to different light / dark intervals. A) NPQ activation measured with a 30-minute treatment with saturating actinic light (800 μmol photons m^−2^ s^−1^); Data were fitted with a logistic function in red. B) Repetition of two 8-minutes (8’) light treatments followed by 10 minutes dark relaxation. C) Repetition of 8 minutes light followed by 90 minutes dark. D) Repetition of two 2-minutes light treatment followed by 10 minutes dark relaxation. Yellow and black boxes indicate light and dark intervals, respectively. In B-D, Asterisks indicate statistically significant differences in NPQ activation during the second light treatment with respect to the first light exposure (tested in the second point after light is switched on, One-way ANOVA, p-value<0.05). Data are expressed as average ± SD of three independent biological replicates.

The impact of zeaxanthin on NPQ can be assessed also by performing multiple NPQ-induction measurements, separated by a dark relaxation (22, 23) (Figure 2). In this protocol, most of NPQ relaxes after the first illumination step, following the dissipation of ΔpH across the thylakoid membrane. NPQ induction during the second illumination, however, is faster because some of the zeaxanthin accumulated is not completely reconverted in the dark interval (Figure 2b). By changing the interval between the two illumination phases it is possible to demonstrate that the pool of zeaxanthin synthesized during the first 8 min of light treatment takes much longer to be completely reconverted into violaxanthin, and its presence accelerates NPQ activation in a second light treatment even if this is separated from the first by 90 min in the dark (Figure 2c). Changing instead the length of light treatment confirmed that zeaxanthin active in NPQ is quickly synthesized but then more slowly reconverted into violaxanthin. As example, only 2 min of illumination are sufficient to accumulate enough zeaxanthin to make NPQ faster in a second measurement after 10 min of dark treatment (Figure 2d).

### Generation of Nannochloropsis strains with altered xanthophyll cycle

To investigate the impact of the xanthophyll cycle on photoprotection mechanisms in *Nannochloropsis*, three independent *vde KO* strains were isolated via homology-directed repair mediated by CRISPR-Cpf1 technology (24, 25) (see Materials and Methods for details). Strains with impaired expression of the *VDE* gene (GENE ID: Naga100041g46) were first selected by phenotypic screening via PAM-Imaging, looking for isolates with reduced NPQ capacity. The insertion of the resistance cassette in the expected genome locus was later validated by PCR (Supplementary figure S2).

Three independent strains overexpressing the *ZEP* gene (*ZEP* OE) were also isolated, after *Nannochloropsis* transformation with a modular vector for effective expression of genes of interest (see supplementary Materials and Methods for details), where the full endogenous *ZEP* gene (Gene ID: Naga100194g2) was cloned. Transformed strains were screened phenotypically by PAM-Imaging, looking for those where NPQ relaxation in the dark was faster than in WT, and RT-PCR was used to validate that they indeed overexpressed the *ZEP* gene (Supplementary figure S3).

These strains were compared to *lhcx1 KO* unable to activate NPQ (26) because of the absence of LHCX1 (Gene ID: Naga100173g12), a protein homologous to LHCX/LHCSR proteins, shown in *Chlamydomonas reinhardtii* and *Phaeodactylum tricornutum* to be essential for NPQ activation (6, 27).

When all strains above were cultivated in flasks at low density and optimal light (i.e. 100 μmol photons m^−2^ s^−1^) for 4 days (see Material and Methods) they showed no differences in growth with respect to the parental strain (Supplementary figure S4). Both *lhcx1* and *vde* KO strains showed a strong reduction of NPQ activation with respect to WT (Figure 3a, b) while the *ZEP* OE strain instead showed a minor reduction in the NPQ activation capacity upon illumination, but also a faster relaxation when the light was switched off with respect to the parental strain (Figure 3c).

**Figure 3.**
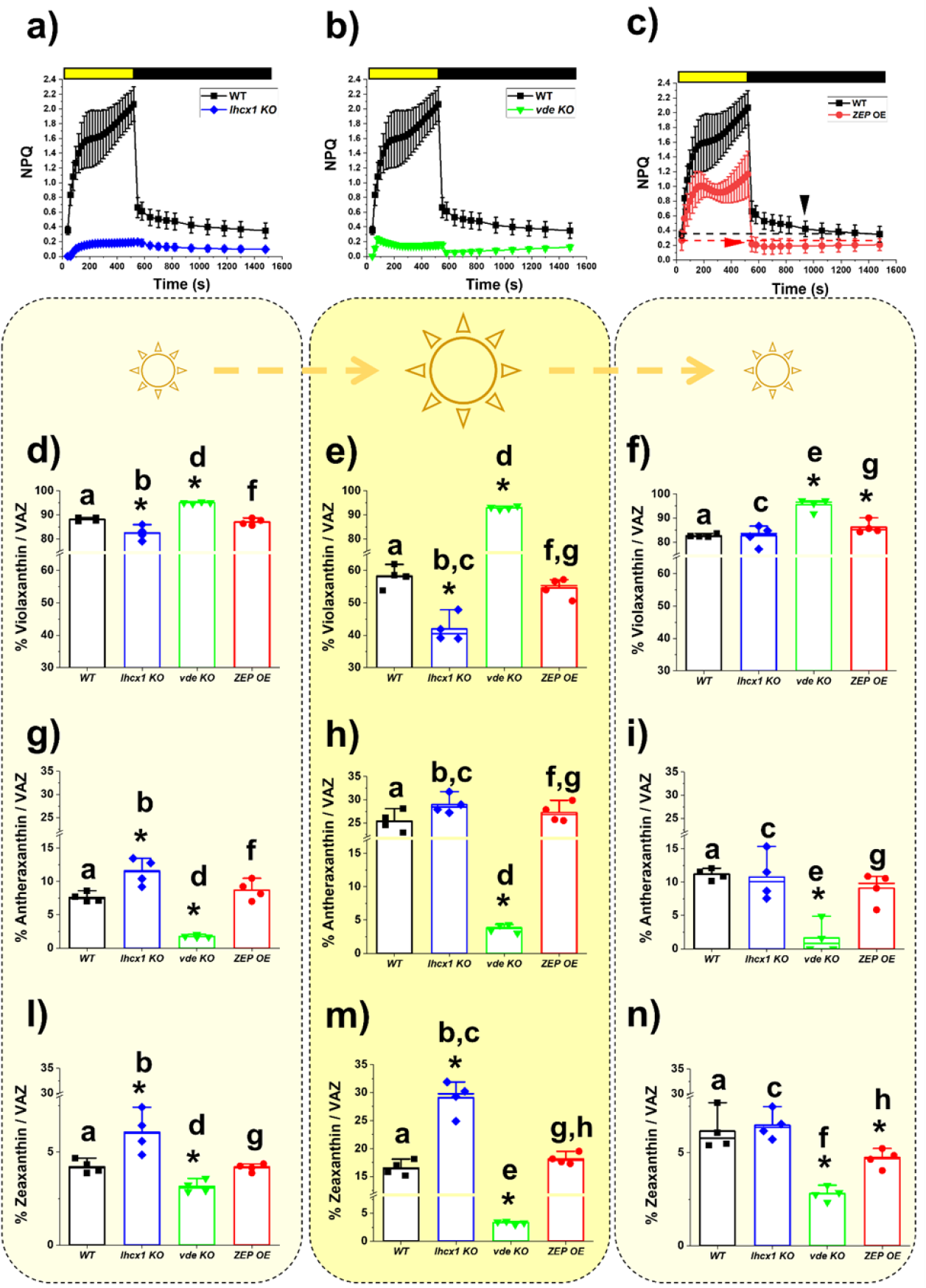
Phenotypic characterization of *Nannochloropsis* strains with altered NPQ response and xanthophyll cycle. NPQ activation and relaxation kinetics for the WT *Nannochloropsis* strain (black squares) upon exposure to saturating light (yellow box) and dark (black box), respectively, compared to the *lhcx1 KO* (blue diamond, a), the *vde KO* (green downward triangle, b) and the ZEP overexpressing strain (red circles, c). The two arrows in panelc) indicate when the NPQ fully relaxes in the two strains. Xanthophylls content after cultivation for 4 days in liquid medium at optimal light (i.e. 100 μmol photons m^−2^ s^−1^) (d,g,l), upon treatment with saturating light (1000 μmol photons m^−2^ s^−1^) for 2 h (e,h,m) and after recovery in optimal light for 1.5 h (f,i,n) for the WT *Nannochloropsis* strain (black), the *lhcx1 KO* (blue), the *vde KO* (green) and the ZEP overexpressing strain (red). Data are expressed as percentage of each xanthophyll molecule over their sum [violaxanthin, (d,e,f); antheraxanthin, (g,h,i) and zeaxanthin (l,m,n); VAZ]. Data are expressed as average ± SD of four independent biological replicates. Asterisks indicate statistically significant differences between each of the mutants and parental strain, in every panel of the figure. Statistically significant differences in the content of each xanthophyll within the same strain, in different conditions (e.g., panels d, e and f) are indicated by the same alphabet letter (One-way ANOVA, p-value<0.05).

In all strains, violaxanthin was the predominant xanthophyll (> 80% VAZ), whilst antheraxanthin and zeaxanthin represent < 10% and < 5% of total VAZ content, respectively. Vaucheriaxanthin and β-carotene were the other major carotenoids detected and they did not show any change in abundance either between genotypes or in the different light conditions tested (Supplementary Table S2), a result consistent with the hypothesis that the genetic modifications of these strains only affected the xanthophyll cycle.

Whilst the *ZEP* OE did not show differences in the content of the three xanthophylls with respect to the parental strain, *lhcx1 KO* showed a reduction in the content of violaxanthin with a corresponding increase of both antheraxanthin and zeaxanthin (Figure 3d, g, l), suggesting that the absence of the LHCX1 protein impacts the xanthophyll cycle as well. The *vde KO* strain showed instead an opposite trend, with an increased accumulation of violaxanthin and a corresponding reduction of the content of antheraxanthin and zeaxanthin with respect to the parental strain (Figure 3d, g, l), suggesting that, in WT cells, VDE in this species has a minor activation even in the relatively low light used here during strain cultivation.

When treated with intense light (1000 μmol photons m^−2^ s^−1^ for 2 h), *lhcx1* KO showed activation of the xanthophyll cycle but, interestingly, the accumulation of zeaxanthin and the corresponding decrease of violaxanthin were higher than in the parental strain (Figure 3e, h, m), suggesting that LHCX1 absence facilitates xanthophyll conversion upon excess light exposure. In *vde* KO, light treatment did not induce any significant change in antheraxanthin and zeaxanthin (Figure 3d, g, l) and, as a result, upon saturating light, the content of violaxanthin was much larger in the *vde* KO than in the WT (Figure 3e, h, m). The *ZEP* OE, instead, did not show major differences in the accumulation of the three xanthophylls upon excess light exposure with respect to WT. This observation can be explained by the possibility that ZEP activity is inhibited under strong illumination by an unknown post-translational mechanism. Alternatively, it is possible that *ZEP* overexpression was not strong enough to overcome endogenous VDE activity upon strong illumination, and thus it did not impact the overall balance of the xanthophyll composition upon prolonged exposure to saturating light (Figure 3e, h, m).

After treatment with saturating light, all strains were then exposed again to optimal light for 1.5 h to monitor xanthophyll cycle relaxation. 1.5 h were not enough to fully relax the xanthophyll cycle in the parental strain (Figure 3f, i, n), as observed before (Figure 1d and 2c). In the same time interval *lhcx1 KO* was instead capable to restore the xanthophyll content measured before excess light exposure (Figure 3f, i, n), demonstrating that the absence of LHCX1 facilitates xanthophyll conversion in both directions. In the same time, *ZEP* OE showed an increased accumulation of violaxanthin and a parallel reduction of zeaxanthin (24% lower with respect to WT) after 1.5 h recovery in optimal light, demonstrating that this strain re-converted zeaxanthin into violaxanthin faster than the parental strain (Figure 3n).

### Impact of xanthophyll cycle on photoprotection

All strains were then tested for their ability to withstand saturating illumination by exposing them for 14 days to 500 μmol photons m^−2^ s^−1^ on agar plates. The *vde* KO showed a strong reduction in growth with respect to the parental strain (Figure 4a, b), demonstrating a major role played by zeaxanthin in photoprotection, whilst no significant differences were detected for the other two strains (Figure 4a, b).

**Figure 4.**
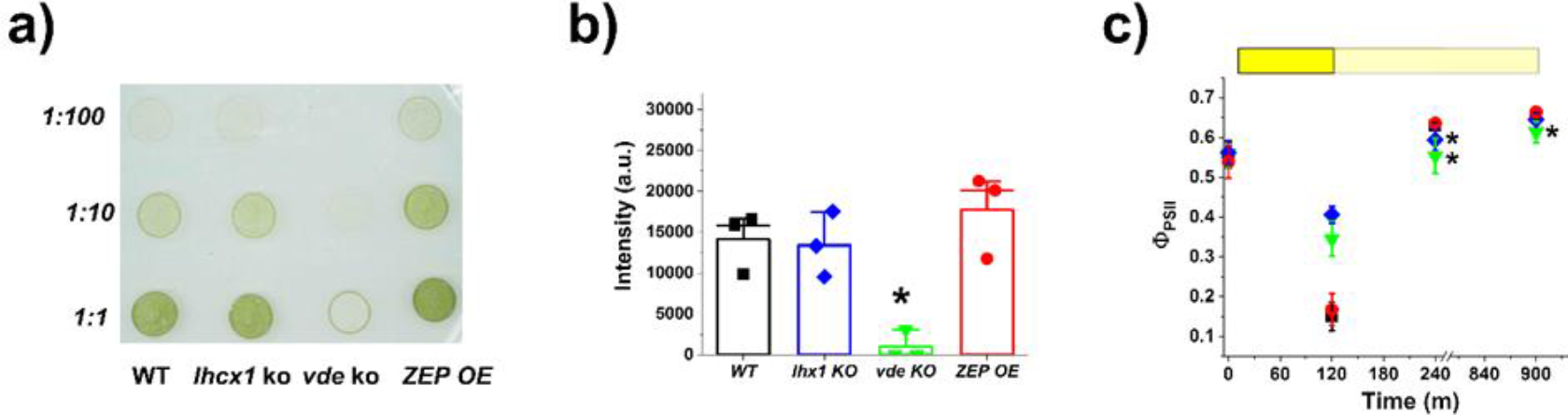
Impact of the xanthophyll cycle on photoprotection. a) Agar plate with spots starting from the same cell concentration for all the strains with different degrees of alteration of the xanthophyll cycle used in this work. Plate was supplemented with 10 mM NaHCO_3_ to avoid carbon limitation and it was grown for 14 days at 500 μmol photons m^−2^ s^−1^. Strain ID is indicated on the bottom, whilst dilution factor on the left. b) Quantification of the intensity of the spots was performed with the software ImageJ (v. 1.52; https://imagej.nih.gov/ij/index.html) and it is here presented for the 1:10 dilution of panel a). c) Photosynthetic efficiency of all the strains used in this work after treatment with saturating light (1000 μmol photons m^−2^ s^−1^) for 2 h (yellow box) and upon recovery in dim light (pale yellow box) for 12 h. WT *Nannochloropsis* strain, black squares; *lhcx1 KO*, blue diamonds; *vde KO*, green downward triangles; ZEP overexpressing strain, red circles. Data are expressed as average ± SD of three independent biological replicates. Asterisks indicate statistically significant differences between mutants and parental strain (One-way ANOVA, p-value<0.05).

To assess the impact of shorter light excess treatments, similar agar plates grown in optimal light (100 μmol photons m^−2^ s^−1^) for 14 days were exposed to saturating light (1000 μmol photons m^−2^ s^−1^) for 2 h while monitoring photosystem II (PSII) quantum yield. All strains showed equal photosynthetic efficiency at the start of the experiment, after growth in optimal light conditions (Figure 4c). Upon exposure to saturating light, there was a strong reduction of photosynthetic efficiency, because of multiple phenomena such as saturation of photosynthetic electron transport, NPQ activation and damage to PSII. Both *lhcx1* and *vde* KO strains showed a smaller reduction than the WT (Figure 4c), explainable by their inability to activate NPQ (Figure 3), while the *ZEP* OE instead showed the same reduction observed in the WT (Figure 4c).

While reoxidation of electron transporters and NPQ relaxation takes a few minutes, PSII photoinhibition takes several hours to be recovered, and this different kinetics can be exploited to distinguish the different contribution to the decrease in photochemical yield observed in Figure 4c. To this aim, cells were allowed to recover under dim light for 12 h, monitoring PSII quantum yield. After 4 h of recovery, *lhcx1 KO* showed a lower PSII quantum yield than the parental strain, suggesting that the mutation led to higher photoinhibition in this strain, although it recovered after 12 h of dim light. *vde KO* showed even larger differences, which were not fully recovered in the time monitored, suggesting that this strain had a larger photosensitivity with respect to the others (Figure 4c).

The importance of both NPQ and the xanthophyll cycle to preserve photosynthetic functionality in over-saturating irradiances was confirmed by monitoring the photosynthetic activity of all strains upon treatment with increasing irradiances (supplementary results and supplementary figure S5). The *lhcx1 KO* and *vde KO* strains *s*howed a faster decrease of qL as the light intensity increased, suggesting their reactions centers were more easily saturated (Supplementary figure S5c), as well as a strong reduction of oxygen evolution upon exposure to increasing light (Supplementary figure S5e), highlighting the importance of NPQ and the xanthophyll cycle to preserve photosynthetic functionality in cells exposed to over-saturating irradiances. *ZEP* OE instead showed a higher photochemical activity than the parental strain at saturating light intensities (Supplementary figure S5c), also confirmed by the slower reduction of PSII activity (Supplementary figure S5d). *ZEP* OE also showed an increase of the photosynthetic electron transport (ETR) that also reached saturation at higher light intensities than the parental strain (Supplementary figure S5b).

### Impact of xanthophyll cycle on biomass productivity in photobioreactors

*Nannochloropsis* strains affected either in NPQ activation or xanthophyll cycle dynamics were cultivated in lab-scale photobioreactors to investigate the impact of photoprotection mechanisms on biomass productivity in industrially relevant conditions. In this setup, microalgae are cultivated in fed-batch mode at high biomass concentration (i.e., 1.5 g · L^-1^, 250 · 10^6^ cells · ml^-1^). Because of the high optical density, the first layers of the cultures are fully exposed to illumination while cells deeper in the volume are in light limitation (28). Environmental complexity is further increased by the culture mixing, causing cells to abruptly move from limiting illumination to full irradiation and *vice versa*.

Cultures were exposed to two irradiances, namely 400 and 1200 µmol photons · m^-2^ · s^-1^, as depicted in supplementary figure S6. Both irradiances are saturating for *Nannochloropsis*, and cells more exposed to illumination thus experience light excess. Because of the culture optical density, however, most of the cells deeper in the culture (approx. > 1 and > 2 cm out of 5 total cm for an incident illumination of 400 and 1200 µmoles photons · m^-2^ · s^-1^, respectively (28)) were still light limited. Cultures were diluted every other day to restore the initial biomass concentration (Supplementary figure S7), and biomass concentration before and after dilution was used to calculate biomass productivity for all the strains investigated (Figure 5a).

**Figure 5.**
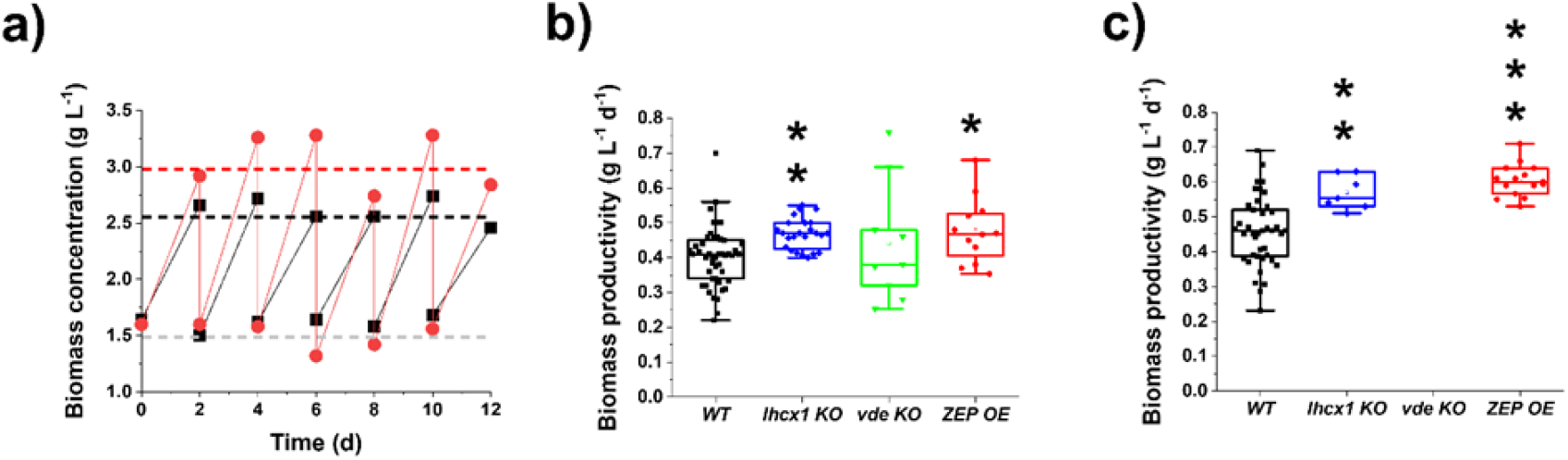
Biomass productivity of *Nannochloropsis* semi-continuous cultures. a) Operational scheme for *Nannochloropsis* semi-continuous cultures. Data were collected before and after dilution to restore the initial biomass concentration of 1.5 g L^-1^, for both WT (black squares) and ZEP over-expressor (red circles). Biomass productivity of *N. gaditana* strains investigated in this work, upon exposure to 400 (b) and 1200 µmol photons · m^-2^ · s^-1^ (c). Asterisks indicate statistically significant differences between the different strains and the WT (One-way ANOVA, * p-value < 0.05; ** p-value < 0.01; *** p-value < 0.001). All strains show a greater biomass productivity at 1200 than at 400 µmol photons · m^-2^ · s^-1^ (One-way ANOVA, p-value < 0.01). Part of the semi-continuous data used to calculate biomass productivity values in b and c are reported in supplementary figure S7.

When exposed to higher irradiance, we observed a reduction in Chl and an increase in Car content for all the strains investigated in this work, indicating activation of an acclimation response (29), but without showing major differences between strains (Supplementary Table S3). Maximal photosynthetic efficiency (ΦPSII) showed a general reduction upon cultivation at stronger irradiance, likely because of some photoinhibition. Maximal photosynthetic efficiency did not show major differences between the strains here investigated, with the exception of *vde* KO (Supplementary Table S4).

With 400 µmol photons · m^-2^ · s^-1^ illumination, *lhcx1* KO and *ZEP* OE showed a higher biomass productivity than the WT, whilst no difference was observed for *vde* KO (Figure 5b). When irradiance increases up to 1200 µmol photons · m^-2^ · s^-1^ all cultures produced more biomass and the difference between the *ZEP* OE and *lhcx1* KO with respect to the parental strain increased, whilst the *vde KO* did not survive (Figure 5c). As shown in Figure S8, *vde KO* was unable to maintain sufficient cell duplication rate and maintain the cell concentration of the culture upon exposure to strong illumination.

In order to confirm the highly different impact of LHCX1 and VDE absence on biomass productivity, analogous mutants impaired in NPQ activation and zeaxanthin biosynthesis (i.e. *lhcx1 KO* and *vde KO*, respectively) from another species of the same genus, *N. oceanica*, were similarly analyzed (30). Also in this case, the *vde KO* strain showed strong sensitivity to high light exposure (Supplementary Figure S8c), while *lhcx1 KO* was fully able to survive high irradiance in dense cultures and showed higher biomass productivity than the WT in these conditions (Supplementary Figure S8b). This confirms that the strong sensitivity of the *vde KO* strain was due to the biological role of zeaxanthin in acclimating to saturating irradiances in *Nannochloropsis*.

### Impact of xanthophyll cycle dynamics on the response to light fluctuations

One major feature of dense cultures in photobioreactors is that microalgae are exposed to inhomogeneous irradiance, and they can suddenly move from excess to limiting light conditions and *vice versa*. To assess in more detail the impact of xanthophyll cycle on response to dynamic light regimes, we simulated the fluctuations of irradiance cells experience in dense cultures and measured the impact on photosynthetic activity, quantified from oxygen evolution using a high sensitivity instrumentation (Figure 6). Light fluctuations were designed to provide, on average, an optimal number of photons for *Nannochloropsis* [i.e. 100 µmol photons · m^-2^ · s^-1^, (26)] but through cycles of saturating and limiting illumination (i.e. 300 and 15 µmol photons · m^-2^ · s^-1^, respectively) for different time frames (i.e. 3 and 7 minutes, respectively) in order to highlight any eventual difference in response to strong illumination or limiting light (Figure 6a).

**Figure 6.**
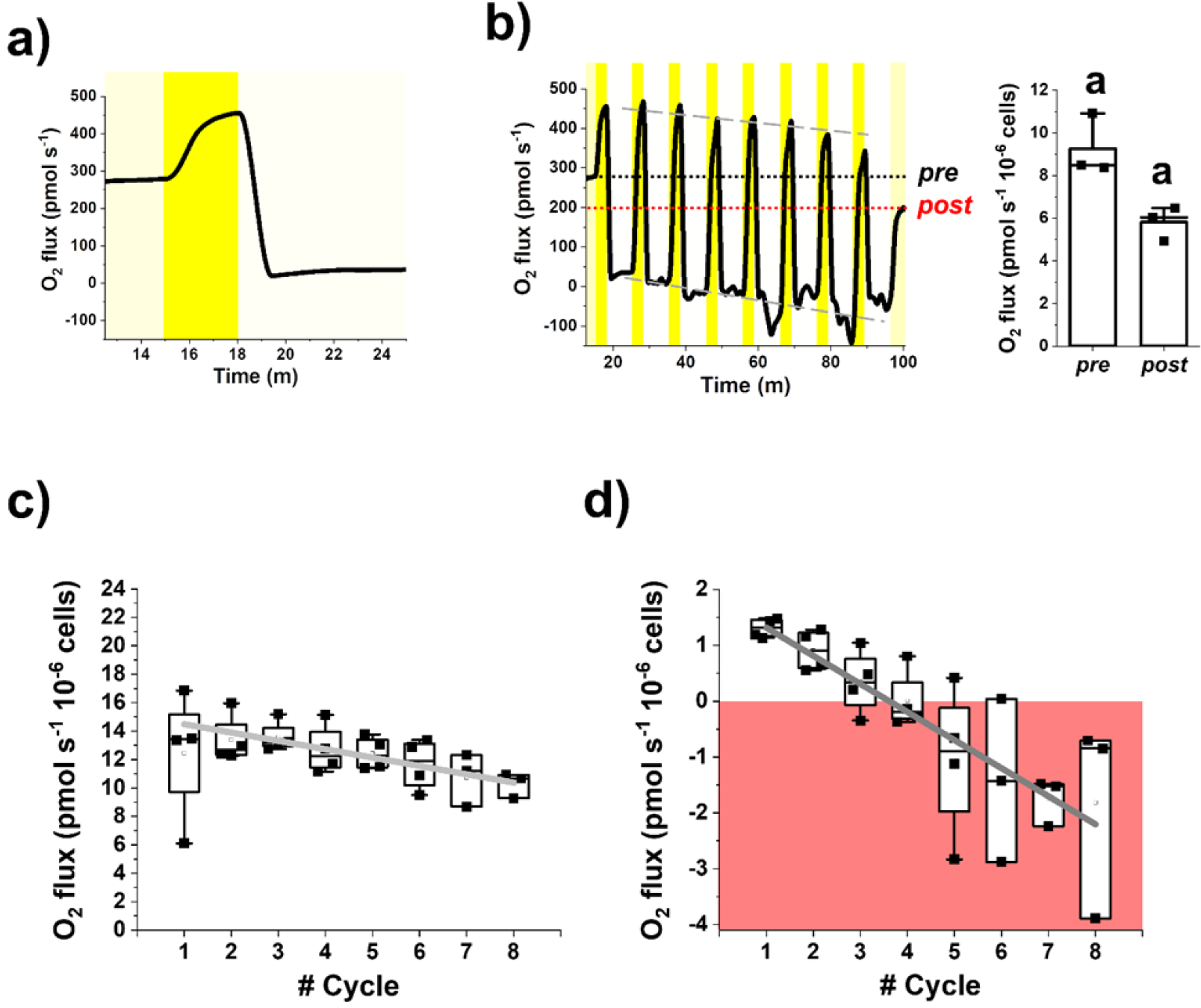
Photosynthetic functionality of WT *Nannochloropsis* in fluctuating light. Oxygen evolution of the WT *Nannochloropsis* strain was measured in 2 ml-samples at a concentration of 100·10^6^ cells/ml (see Materials and Methods for details). a) we designed a method to treat cells with a light fluctuation protocol where they were first exposed to optimal light at 100 µmol photons m^-2^ · s^-1^ (yellow box) until a steady photosynthetic activity was reached, then to 300 µmos photons · m^-2^ · s^-1^ (dark yellow box) and 15 µmol photons · m^-2^ · s^-1^ (light yellow box) for 3 and 7 minutes, respectively. Irradiance and time of exposure were set so to provide cells with an optimal number of photons, corresponding to 100 µmol photons · m^-2^ · s^-1^. b) the two phases at 300 and 15 µmol photons · m^-2^ · s^-1^ were repeated 8 times and after cells were returned to an optimal irradiance of 100 µmol photons · m^-2^ · s^-1^. Black and red dot lines indicate the oxygen flux at 100 µmol photons m^-2^ · s^-1^ before (pre) and after (post) light fluctuation, respectively. Grey dashed lines instead indicate the trend of oxygen flux over the fluctuation cycles. The oxygen evolution activity pre- and post-light fluctuation were compared to measure the impact of light fluctuation on photosynthetic activity (right plot in panel b). The same alphabet letter indicates statically significant differences between oxygen evolution values at 100 µmol photons · m^-2^ · s^-1^, before and after light fluctuation (One-way ANOVA, p-value<0.05). Oxygen evolution activity of WT cells at 300 µmol photons · m^-2^ s^-1^ c) and 15 µmol photons · m^-2^ · s^-1^ (d) over the number of fluctuation cycles. Data at a specific light intensity come from the average oxygen evolution rate measured over 20 seconds of the trace in a). In d) the area where oxygen consumption via respiration is higher than oxygen evolved via photosynthesis is highlighted by a red box. At both irradiances, photosynthetic activity significantly drops (slope is significantly different from zero, one-way ANOVA, p-value < 0.05) over the fluctuation cycles, according to the following linear functions: y = (15.07 ± 0.35) – (0.59 ± 0.06) x, Pearson’s R: -0.97, R-Square: 0.94 (c); y = (1.82 ± 0.045) – (0.5 ± 0.01) x, Pearson’s R: -0.99, R-Square: 0.99 (d). Data are expressed as average ± SD of four independent biological replicates.

O2 evolution in WT cells changed following the light irradiance dynamics, as expected (Figure 6a and supplementary table S5). Cells were first exposed to a constant optimal light intensity at 100 µmol photons · m^-2^ · s^-1^ to reach a steady photosynthetic activity (9.3 ± 1.4 pmol O2 s^-1^ 10^−6^ cells). When light increased to 300 µmol photons · m^-2^ · s^-1^, photosynthetic activity followed, reaching a new steady state after approx. 2 min (12.45 ± 4.5 pmol O2 s^-1^ 10^−6^ cells). When light decreased to 15 µmol photons · m^-2^ · s^-1^, photosynthetic activity decreased to reach a lower steady oxygen evolution rate after approx. 4 min (1.31 ± 0.17 pmol O2 s^-1^ 10^−6^ cells, Supplementary table S5).

The same light fluctuation was then repeated 8 times covering a total of 80 minutes, followed by another exposure at optimal constant light at 100 µmol photons · m^-2^ · s^-1^ (Figure 6b). The repetition of light fluctuations had a clear effect on *Nannochloropsis* photosynthetic activity. The oxygen evolution activity of the WT at steady 100 µmol photons · m^-2^ · s^-1^ illumination after the fluctuation treatment was significantly reduced to 5.8 ± 0.8 pmol O2 s^-1^ 10^−6^ cells, 37% lower than before (Figure 6b). Consistently, the trace in Figure 6b suggested that also oxygen evolution activities at 300 and 15 µmol photons · m^-2^ · s^-1^ progressively decreased with each fluctuation cycle, as confirmed when these trends were analysed in detail, showing a significant linear decay (Figure 6c and d, respectively). Clearly these data suggest that light fluctuations caused a decrease in photosynthetic activity, because of the activation of photo-regulatory mechanisms and photoinhibition.

The reduction of photosynthetic rates observed at 15 µmol photons · m^-2^ · s^-1^ is relatively larger than the one observed at 300 µmol photons · m^-2^ · s^-1^ (Figure 6d and 6c, respectively). Even more importantly, at low illumination the activity became negative, meaning that in these cells photosynthesis is not able to compensate for respiration (Figure 6d). These data are particularly informative on the behavior of microalgae cells in dense cultures of industrial systems, where cells are exposed to continuous light fluctuations and a large fraction of the culture volume is light limited (18), and suggest that these cells might indeed have negative photosynthetic activity, thus curbing overall photon-to-biomass conversion efficiency and biomass productivity.

The strains affected in photoprotection and the xanthophyll cycle were also exposed to a similar light profile. At 100 µmol photons · m^-2^ · s^-1^ constant illumination, *vde KO* and *ZEP OE* showed the same photosynthetic activity of the WT, whilst *lhcx1 KO* instead showed a significant reduction (Supplementary Table S5). After exposure to light fluctuations, *vde KO* showed a significant reduction of photosynthetic activity at 100 µmol photons · m^-2^ · s^-1^ (32%), similar to WT. On the contrary, the photosynthetic activities of both *lhcx1 KO* and *ZEP OE* were not affected (Figure 7 a, c and e). The phenotype of *lhcx1 KO* suggests that the reduction of oxygen evolution activity upon exposure to light fluctuations observed in the WT is due to NPQ activation. On the other hand, *vde KO* showed a decrease too, likely attributable to the strong photosensitivity of this strain.

**Figure 7.**
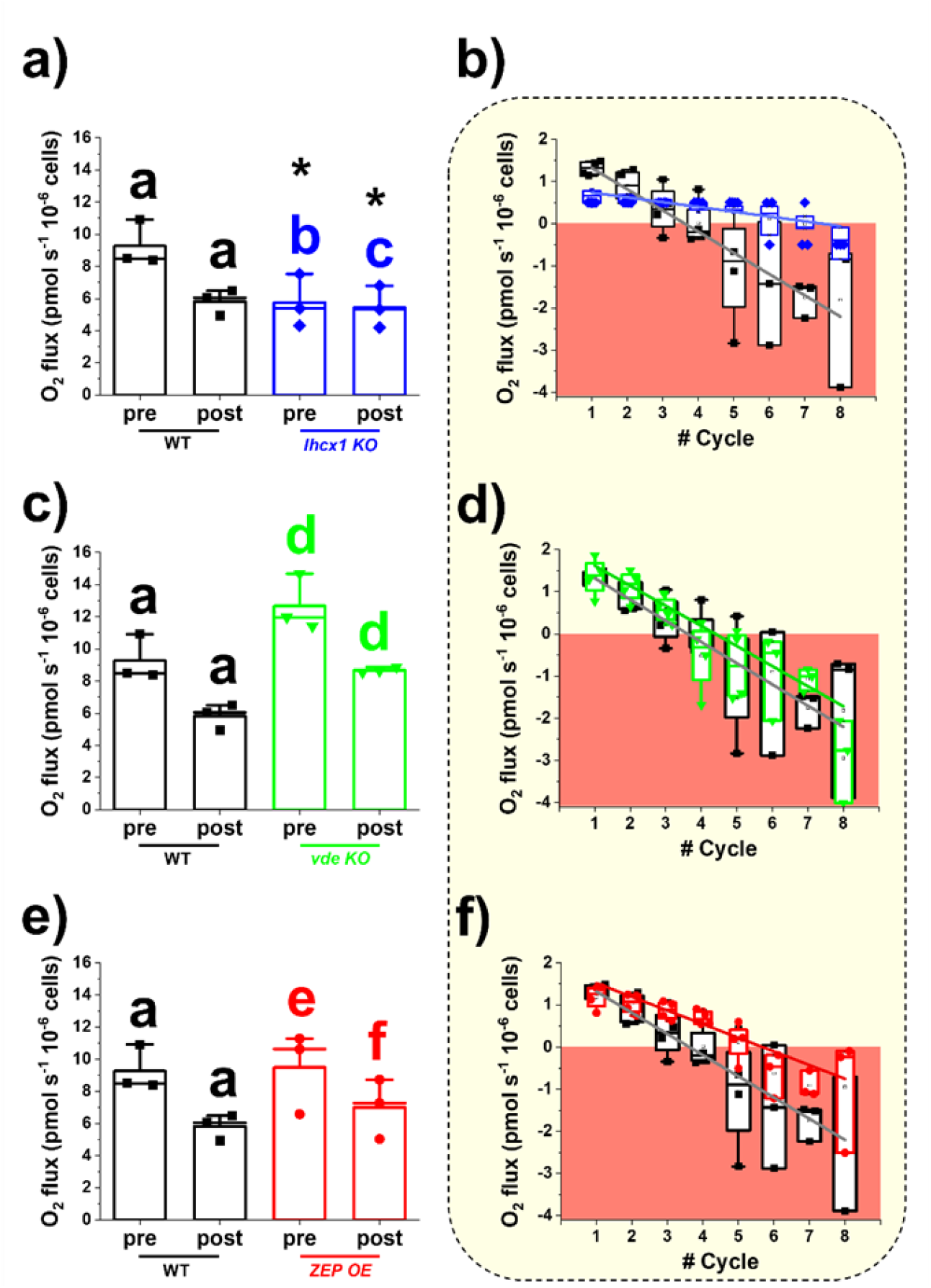
Photosynthetic functionality of strains affected in NPQ and xanthophylls cycle dynamics in fluctuating light. Photosynthetic functionality is expressed as oxygen evolution activity and was measured in the conditions described in Figure 6. Oxygen evolution activity of *lhcx1 KO* (a), *vde KO* (c) and *ZEP OE* (e) before (pre) and after (post) the light fluctuation treatment of Figure 6, compared to the WT. The same alphabet letter indicates statically significant differences between oxygen evolution values at 100 µmol photons · m^-2^ · s^-1^, before and after light fluctuation within the same strain, whilst asterisks indicate statistically significant differences between mutants and WT (One-way ANOVA, p-value<0.05). Oxygen evolution activity for *lhcx1 KO* (blue diamonds, b), *vde KO* (green downward triangles, d) and *ZEP OE* (red circles, f) cells at 15 µmol photons · m^-2^ · s^-1^ (light yellow box) over the number of fluctuation cycles compared to the WT (black squares). The area where oxygen consumption via respiration is higher than oxygen evolved via photosynthesis is highlighted by a red box. The linear oxygen evolution trend over the cycles of fluctuation has been mathematically described by the functions reported in Supplementary table S6. Data are expressed as average ± SD of four independent biological replicates.

All mutant strains showed a significant reduction of the oxygen evolution activity at 15 µmol photons · m^-2^ · s^-1^ over the cycles of fluctuations, as observed in the WT (Figure 7b, d, f and supplementary table S6). Both *lhcx1 KO* and *ZEP OE* showed a smaller reduction over the cycles of fluctuations and oxygen evolution activity at 15 µmol photons · m^-2^ · s^-1^ became negative after 5 cycles, different from WT and *vde KO* where negative values were reached only after 3 cycles of fluctuation (Figure 7c, f and i). After the treatment, oxygen evolution at 15 µmol photons · m^-2^ · s^-1^ was -0.44 ± 0.37 pmol O2 s^-1^ 10^−6^ cells for *lhcx1 KO* and -0.95 ± 1.35 pmol O2 s^-1^ 10^−6^ cells for *ZEP OE*, while -1.81 ± 1.7 pmol O2 s^-1^ 10^−6^ cells for WT and -2.9 ± 0.9 pmol O2 s^-1^ 10^−6^ cells for *vde KO* (Figure 7).

## Discussion

### Biological role of zeaxanthin in Nannochloropsis

*Nannochloropsis gaditana* cells upon exposure to excess light show the ability to convert violaxanthin into zeaxanthin (Figure 1), as in many other photosynthetic eukaryotes (17). *Nannochloropsis* has a peculiar pigment composition with violaxanthin being the most abundant carotenoid in this species, accounting for approx. 50% of the total (31–33). Likely because of this large reservoir of substrate, in contrast to plants and other microalgae (34, 35), zeaxanthin synthesis in *Nannochloropsis* continues even upon prolonged exposure to extreme irradiances with no visible saturation (Figure 1). Considering the light intensities tested in this work, which went well beyond physiologically relevant conditions, our results also suggest that zeaxanthin synthesis is unlikely to ever reach saturation in the natural environment, meaning that *Nannochloropsis* cells are capable of additional zeaxanthin synthesis whenever needed even if they have already been exposed to strong illumination.

The large capacity of zeaxanthin synthesis is accompanied by a strong impact of this pigment on the protection of the photosynthetic apparatus. The phenotype of both *vde* KO and WT cells treated with the VDE inhibitor DTT demonstrate that zeaxanthin synthesis has a major impact on NPQ in *Nannochloropsis* (Figure 3b and supplementary figure S1), as also observed in (30).

Zeaxanthin synthesis impacts NPQ from the first few seconds of illumination (Figure 2), while HPLC analysis shows that a few minutes of illumination are needed before detecting a significant accumulation of molecules (Figure 1). This observation suggests that a small number of zeaxanthin molecules can activate NPQ in a few seconds after an increase of illumination, likely by associating to specific binding sites in light-harvesting complexes. Considering that also *lhcx1* KO strain shows a major decrease in NPQ capacity, and that its full activation requires the presence of both zeaxanthin and LHCX1, it is likely that zeaxanthin activity in NPQ requires its association to the LHCX1 protein in *N. gaditana*, as previously suggested for *N. oceanica* (30). Similarly, in diatoms NPQ is provided by a concerted action between LHCX proteins and diatoxanthin (36), a xanthophyll molecule part of the diadinoxanthin-diatoxanthin cycle, which is analogous to the VAZ cycle observed in *Nannochloropsis* (37). LHCX1 is the main NPQ effector also in diatoms, although additional LHCX proteins, namely LHCX2 and LHCX3, are involved when cells are exposed to prolonged high light, providing flexibility of quenching site but most likely with a similar mechanism (36, 38, 39).

Pigment data of the *lhcx1 KO* strain also show that the absence of LHCX1 has a measurable impact on the xanthophyll cycle dynamics with a larger accumulation of zeaxanthin than in WT, but also a faster conversion back to violaxanthin. This can be explained knowing that a large fraction of violaxanthin is bound to antenna proteins and it needs to be released into the thylakoid membrane to be converted into zeaxanthin. This exchange from antenna proteins limits the rate of xanthophyll conversion, as demonstrated in plants (40). *lhcx1 KO* is depleted of one of the most abundant antenna proteins in *Nannochloropsis* (41), and this is likely to accelerate zeaxanthin synthesis and degradation because of a larger presence of carotenoids not bound to antenna proteins, but free in the thylakoid membranes and thus more available to VDE.

In *N. gaditana*, even though NPQ slowly continues to increase after 10 min induction, suggesting the presence of a qZ-type contribution associated with the progressive accumulation of zeaxanthin, the largest fraction of NPQ capacity reaches saturation in this time frame (Figure 2). Since zeaxanthin synthesis continues much longer without showing signs of saturation (Figure 1), this suggests that it is rather the influence of zeaxanthin molecules on NPQ that is slowing down, likely because of saturation of the potential binding sites for zeaxanthin in LHCX1. A second pool of zeaxanthin molecules continues to be synthesized upon prolonged exposure to strong light, but it does not contribute to NPQ and likely plays other roles in photoprotection such as direct scavenging of Chl triplets and ROS (9).

While the zeaxanthin molecules active in NPQ are quickly synthesised, their impact on NPQ remains for a prolonged time. This is evidenced by the fact that NPQ induction kinetics are faster if cells have already been exposed to a previous light treatment (Figure 2). This effect is already visible after exposing cells to light for 2 min and it is still detectable after a 90-min dark relaxation, demonstrating that this time is not sufficient to re-convert all zeaxanthin synthesized in 8 min illumination (Figure 2). This effect can be modulated by overexpressing ZEP since cells are faster in re-converting zeaxanthin into violaxanthin during the 90-min dark relaxation, as demonstrated by the reduction in NPQ induction during the second kinetic with respect to the parental strain (Supplementary figure S9), supporting the HPLC data of Figure 3.

### Zeaxanthin plays an essential photoprotective role in Nannochloropsis, beyond NPQ

Both *vde KO* and *lhxc1* KO strains show sensitivity to saturating illumination, supporting the role of NPQ on protection of *Nannochloropsis* from light stress (Figure 4). When cells are cultivated in dense cultures, however, the results between the two genotypes are very different. In this context some cells are exposed to full illumination, while the others, because of shading, are in limiting light or even dark (28). In the experimental system employed here, approx. 60% of incident radiation is absorbed by the 1^st^ cm of culture depth (18). If the culture is exposed to a strong external illumination (1200 µmol photons · m^-2^ · s^-1^), *vde* KO cells show a clear decrease in maximum quantum yield of PSII (Supplementary Table S4), suggesting that more exposed cells are extensively damaged by illumination. This damage cannot even be counterbalanced by cells deeper in the culture volume and eventually it impairs the growth of the whole culture under strong illumination (Supplementary figure S8).

The inability of the *vde* KO strain to grow at higher illumination depends on its stronger photosensitivity as a consequence of the absence of both the NPQ response and the activation of the xanthophyll cycle upon exposure to saturating irradiance, as demonstrated in Figures 3, 4 and 5. While both *vde KO* and *lhcx1* KO strains are similarly defective in NPQ (Figure 3), the latter retains growth under strong illumination, clearly demonstrating that the impact of zeaxanthin biosynthesis on photoprotection goes well beyond its role in enhancing NPQ and that its ability to increase scavenging of Chl triplets and ROS (9, 42) is essential even in dense cultures.

### Xanthophyll cycle dynamics has a major impact on microalgae biomass productivity in photobioreactor

Microalgae at industrial scale are cultivated at high concentration to maximize biomass productivity. Such dense cultures are also continuously mixed to maximize the exposure of cells to incident light and avoid nutrient and carbon limitation, causing cells to suddenly move between limiting and excess illumination, further increasing the complexity of the light environment. In these environmental conditions, more exposed cells need photoprotection mechanisms to withstand strong illumination, but the same mechanisms become detrimental for productivity once the cells move to light limitation of deeper layers. The trade-off between photoprotection and photochemical efficiency, which must be balanced by all photosynthetic organisms (3), is thus particularly challenging in such a complex and dynamic environmental context. It is not surprising that strategies for the optimization of photosynthetic productivity have generated mixed results so far (43, 44), with the only reasonable conclusion being that the complexity of the natural and artificial changes experienced by microalgae during industrial cultivation has a major influence on productivity that cannot be underestimated (45).

Strains with altered xanthophyll cycle analysed in this work demonstrate that an efficient photoprotection is essential for microalgae fitness in dense cultures to ensure growth under full sunlight, as shown by the strong photosensitivity of *vde* KO. On the other hand, we observed that *lhxc1* KO in dense cultures shows a positive impact on biomass productivity. This strain differs from WT because of its reduction in NPQ activation, but these cells also have a reduced PSII antenna size and Chl content per cell (46) and a higher zeaxanthin content, observed in this work. Mathematical models suggest that the reduction in Chl content per cell should have the largest impact in improving biomass productivity (46), but it is also possible that the higher zeaxanthin content observed in *lhcx1* KO can compensate for any eventual extra damage due to NPQ inactivation.

Energy losses due to natural kinetics of photoprotection can be detrimental for productivity, and accelerating zeaxanthin conversion to violaxanthin can be advantageous in this context. In this work we also simulated the light fluctuation experienced by microalgae in dense cultures of industrial systems (Figure 6b) as a consequence of mixing and observed that WT cells showed a substantial reduction of photosynthetic functionality in light limitation after only a few fluctuation cycles (Figure 6d). This decrease could be due to multiple phenomena, such as the activation of photoprotection or photoinhibition. The *lhcx1 KO* strain does not show the same reduction of WT, suggesting that NPQ is the major factor responsible for the loss of activity observed in the parental strain in dense cultures. On the other hand, the *vde KO* strain showed an even larger reduction of photosynthetic functionality in light limitation (Figure 7f), suggesting that photoinhibition can also play a major role.

In the case of the *ZEP* OE, cells maintain the ability to activate NPQ but also have faster recovery, suggesting that increasing the rate of violaxanthin biosynthesis alone has a beneficial effect on productivity. This is achieved because cells still maintain the ability to synthesize zeaxanthin when needed for photoprotection (Figure 3), but they also have a faster re-conversion rate to violaxanthin when light becomes limiting. This likely provides an advantage when cells move from external to internal, light-limited positions in the dense culture where they remove zeaxanthin faster and can therefore channel more energy towards photochemistry.

It is also worth noting that light-limited layers represent the major fraction of the volume in dense cultures of industrial systems (18), suggesting that an improved photochemical activity in these layers is likely to provide the greatest impact on productivity. This is consistent with the observation that *lhcx1 KO* and *ZEP* OE, the two strains that show the smaller reduction in photosynthetic activity upon exposure to light fluctuations, also showed an increase in biomass productivity in dense cultures (Figure 5). This suggests the optimization of the xanthophyll cycle is a valuable strategy in photosynthesis engineering, yet a fine tuning is preferable to an indiscriminate activation, likely because in the latter case the improvement in cell fitness cannot fully compensate the metabolic burden of a hyper-active xanthophyll cycle.

### Optimization of xanthophyll dynamics in microalgae vs plants

The genetic modification of NPQ and xanthophyll cycle has already been demonstrated to be effective to improve biomass productivity in crop plants in the field (14, 15). In our current work, effects are observed in *Nannochloropsis* by overexpressing only ZEP. It is in fact worth mentioning that VDE activity remains strong in the *ZEP* OE strain, such that it is still fully capable of producing zeaxanthin upon excess light exposure. This is likely also connected with a high violaxanthin content of *N. gaditana* with respect to plants, suggesting that this organism likely also has high endogenous VDE activity.

However, when metabolic engineering is applied to photosynthesis, the complexity of the environmental conditions of the intended cultivation system should also be considered, as well as the physiology of the species targeted for improvement. For instance, in plants of *Nicotiana benthamiana, Arabidopsis thaliana* and *Solanum tuberosum*, VDE, ZEP and PSBS overexpression did not show the same effects (47, 48), indicating that species-specific physiological or morphological features are highly influential on the homeostasis of the photosynthetic metabolism. In the environment of photobioreactors, most of the culture is light limited, while only a small layer of cells is exposed to full sunlight. The design of photobioreactors, as well as operational conditions (e.g. culture concentration) strongly affect the percentage of cells that are in light-limiting conditions or excess light, affecting the optimal balance between photoprotection and photochemical efficiency. Culture mixing is also expected to play a major role on this balance. It is then worth noting that the complexity of the natural and artificial changes experienced by microalgae in dense cultures of industrial systems is likely to prevent the identification of ideal strains more productive in all operational conditions, suggesting that photosynthesis optimization efforts should be tuned to the specific operational conditions in use.

## Materials and Methods

### Isolation of vde KO strain in Nannochloropsis

*Nannochloropsis vde KO* mutant strain was isolated via homology directed repair mediated by CRISPR-Cpf1 technology, using recombinant ribonucleoproteins (RNPs). The construct to drive homology repair was designed to contain a cassette conferring resistance to Zeocin (49), flanked on both sides by 1.5 kb genomic regions homologous to the 5’ and 3’ of the *VDE* gene of *Nannochloropsis* (Gene ID: Naga100041g46). The homology repair cassette was then excised from the holding vector and used to transform *Nannochloropsis* according to (49). Prior to transformation, 4 µl of three synthetic RNPs, assembled using recombinant Cpf1 and synthetic sgRNAs (IDT Technologies, USA) in an equimolar ratio (6 µM), at RT for 20 min, were added to the sample to drive three independent events of site-directed double-strand cleavage in the *VDE* gene. sgRNAs sequences used in this work from 5’-3’: 1. gaccaccgcgcgggtgacggcgg; 2. cgtgcagggcgaccggctctacg; 3. gcgaggtcgccgggtttctggtt.

### Strains, cultivation conditions and growth monitoring

*Strains*. In this work we used two species: *Nannochloropsis gaditana* and *Nannochloropsis oceanica*. All strains used in this work are summarized in Table 1.

**Table 1.** *Nannochloropsis* strains used in this work.

| Species | Strain | Reference |
| --- | --- | --- |
| *N. gaditana* | WT | CCAP |
|  | *lhcx1 KO* | (26, 49) |
|  | *ZEP OE* | This work |
|  | *vde KO* | This work |
| *N. oceanica* | WT | CCMP |
|  | *lhcx1 KO* | (30) |
|  | *vde KO* | (30) |

*N. gaditana*, strain CCAP 849/5 was purchased from the Culture Collection of Algae and Protozoa (CCAP). *N. gaditana lhcx1 KO* was previously obtained by insertional mutagenesis (26, 49). *N. gaditana* strains *vde KO* and the *ZEP* over-expressor were generated in this work, the former via CRISPR-Cpf1 whilst the latter after transformation with a cassette conferring resistance to zeocin (49) flanking another one expressing the coding sequence of the endogenous *ZEP* gene (Gene ID: Naga100194g2).

*N. oceanica* strain CCMP 1779 was purchased from the Culture Collection of Marine Phytoplankton (CCMP) and both *vde KO* and *lhcx1 KO* strains were previously generated (30).

*Cultivation conditions*. All microalgae strains of this work were maintained in F/2 solid media, with 32 g/L sea salts (Sigma Aldrich), 40 mM Tris-HCl (pH 8), Guillard’s (F/2) marine water enrichment solution (Sigma Aldrich), 1% agar (Duchefa Biochemie). Cells were pre-cultured in sterile F/2 liquid media in Erlenmeyer flasks irradiated with 100 µmol photons m^-2^ s^-1^, 100 rpm agitation, at 22 ± 1 °C in a growth chamber.

In order to investigate xanthophyll accumulation dynamics in *N. gaditana*, cells were grown in a Multicultivator MC 1000-OD system (Photon Systems Instruments, Czech Republic) in liquid F/2 starting from 10·10^6^ cells/ml, where constant air bubbling provides mixing and additional CO_2_. Temperature was kept at 22 ± 1 °C and different light intensities were provided using an array of white LEDs.

Liquid cultures for phenotypic characterization and monitoring of photosynthetic functionality of the strains investigated in this work started from pre-cultures grown in conditions described above. Cells were washed twice in fresh F/2 before starting growth curves from 5·10^6^ cells/ml in F/2 supplemented with 10 mM NaHCO_3_ to avoid carbon limitation, in Erlenmeyer flasks irradiated with 100 µmol photons m^-2^ s^-1^, 100 rpm agitation, at 22 ± 1 °C in a growth chamber.

Semi-continuous growth was performed at 22 ± 1 °C in 5-cm Drechsel bottles, illuminated from one side, with 250 ml working volume. Mixing and carbon source was provided through the insufflation of air enriched with 5% CO_2_ (v/v) at 1 L h^-1^. In this case, F/2 growth media was enriched with added nitrogen, phosphate and iron sources (0.75 g L^-1^ NaNO3, 0.05 g L^-1^ NaH2PO4 and 0.0063 g L^-1^ FeCl3 · 6H2O final concentrations). Light was provided through cool white fluorescent lamps. Illumination rate was determined using the LI-250A photometer (Heinz-Walz, Effeltrich, Germany). Cultures were maintained in a semi-continuous mode diluting the culture every other day, as described in (28). Cell concentrations was monitored before and after dilution with an automatic cell counter (Cellometer Auto X4, Cell Counter, Nexcelom). All experiments were conducted maintaining cell concentration at 250 · 10^6^ cells · ml^-1^ (∼ 1.5 g L^-1^) and exposing cultures to different light conditions: 400 and 1200 µmol photons m^-2^ s^-1^ (Supplementary figure S1).

*Biomass productivity*. Biomass productivity of semi-continuous cultures was estimated monitoring the dry weight of the culture in semi-continuous mode before and after dilution. Cultures were filtered using 0.45 μm filters, dried at 60 °C for 24 h and weighed (28).

*High light treatment*. High light treatments were performed using a LED Light Source SL 3500 (Photon Systems Instruments, Brno, Czech Republic). Cells were mixed in a thin cylinder placed in a water bath in order to get a homogeneous irradiance and a constant temperature.

### Pigment extraction

Total pigments were extracted in the dark using 1:1 ratio of 100% N, N-dimethylformamide (Sigma Aldrich), for at least 24 h in the dark at 4 °C (50). Absorption spectra were registered between 350 and 750 nm using Cary 100 spectrophotometer (Agilent Technologies) to determine pigment concentration using specific extinction coefficients (50). Absorption values at 664 and 480 nm were used to calculate the concentrations of chlorophyll a and total carotenoids, respectively.

The content of individual carotenoids was determined after extraction with 80% acetone preceded by mechanical lysis using a Mini Bead Beater (Biospec Products) in the presence of glass beads (150–212 μm diameter, Sigma Aldrich), using a high-pressure liquid chromatography (HPLC) as previously described (51). The HPLC system consisted of a 139 reversed-phase column (5μm particle size; 25×0.4 cm; 250/4 RP 18 Lichrocart, Darmstadt, Germany) and a diode-array detector to record the absorbance spectra (1100 series, Agilent, 141 Waldbronn, Germany). The peaks of each sample were identified through the retention time and absorption spectrum (52). The vaucheriaxanthin absorption factor was estimated by correcting that of violaxanthin for their different absorption at 440 nm.

### Fluorescence measurements for the monitoring of NPQ and photosynthetic functionality

The estimation of photosynthetic parameters was performed by measuring *in vivo* Chl fluorescence using a Dual PAM-100 fluorimeter (Heinz-Walz, Effeltrich, Germany). Samples were dark-adapted for 20 min, then exposed at 850 µmol photons m^-2^s^-1^ (actinic light) and dark for different time frames to assess the activation and relaxation trends of NPQ in double kinetics (Figure 2), respectively. For the phenotypic characterization of the strains investigated in this work, samples were instead exposed to 2000 µmol photons m^-2^s^-1^ (actinic light) for 8 min and to dark for 15 min (Figure 3). Photosynthetic functionality was monitored by treating samples at increasing irradiances of actinic light (Supplementary Figure S5). In all protocols, saturating pulses and measuring light were set at 6000 and 42 µmol photons m^-2^s^-1^, respectively. Maximum quantum yield of PSII (ΦPSII), quantum yield of PSII in light-treated samples (Φ’PSII), qL and NPQ were calculated according to (53, 54).

### Oxygen evolution

Oxygen evolution was measured with a start-up O2K-Respirometer (NextGen-O2k and the PB-Module from Oroboros Instruments GmbH, Austria) in 2 ml samples at a concentration of 100·10^6^ cells/ml in F/2 supplemented with 5 mM NaHCO_3_ to avoid carbon limitation, in measuring chambers magnetically stirred at 750 rpm and with a frequency of 2 seconds. The light source was a blue LED with an emission peak at 451 nm (Osram Oslon). Instrument calibration was performed in the same medium and samples were dark adapted for 10 min to assess respiration rate before starting the measurements of the oxygen flux at increasing irradiances. Respiration and photosynthesis rates were measured with the software DatLab 7.4.0.4.

### Statistical analysis

Descriptive statistical analysis was applied for all the data presented in this work. Statistical significance was assessed by one-way analysis of variance (One-way ANOVA) using OriginPro 2018b (v. 9.55) (http://www.originlab.com/). Samples size was at least >4 for all the measurements collected in this work and for biomass productivity it reached >10 data points for all strains investigated.

## Acknowledgments

Authors acknowledge the support from Antoni Mateau Vera-Vives, Katarzyna Krawczyk, Andrea Meneghesso, Andrea Cailotto and Matteo Scarsini for preliminary experiments.

TM acknowledges the support from European Union H2020 Project 862087-GAIN4CROPS. DL and KKN were supported by the U.S. Department of Energy, Office of Science, Basic Energy Sciences, Chemical Sciences, Geosciences, and Biosciences Division under field work proposal 449B. KKN is an investigator of the Howard Hughes Medical Institute.

## References

1. C. de Vargas, et al., Ocean plankton. Eukaryotic plankton diversity in the sunlit ocean. Science 348, 1261605 (2015).

2. D. R. Ort, et al., Redesigning photosynthesis to sustainably meet global food and bioenergy demand. Proc Natl Acad Sci U S A 112, 8529–36 (2015).

3. A. Alboresi, M. Storti, T. Morosinotto, Balancing protection and efficiency in the regulation of photosynthetic electron transport across plant evolution. New Phytologist 221, 105–109 (2019).

4. Z. Li, S. Wakao, B. B. Fischer, K. K. Niyogi, Sensing and responding to excess light. Annu Rev Plant Biol 60, 239–60 (2009).

5. X. P. Li, et al., A pigment-binding protein essential for regulation of photosynthetic light harvesting. Nature 403, 391–5 (2000).

6. G. Peers, et al., An ancient light-harvesting protein is critical for the regulation of algal photosynthesis. Nature 462, 518–21 (2009).

7. R. C. Bugos, H. Y. Yamamoto, Molecular cloning of violaxanthin de-epoxidase from romaine lettuce and expression in Escherichia coli. Proceedings of the National Academy of Sciences 93, 6320–6325 (1996).

8. P. Arnoux, T. Morosinotto, G. Saga, R. Bassi, D. Pignol, A structural basis for the pH-dependent xanthophyll cycle in Arabidopsis thaliana. Plant Cell 21, 2036–44 (2009).

9. M. Havaux, L. Dall’osto, R. Bassi, Zeaxanthin has enhanced antioxidant capacity with respect to all other xanthophylls in Arabidopsis leaves and functions independent of binding to PSII antennae. Plant Physiol 145, 1506–20 (2007).

10. B. Demmig-Adams, J. J. Stewart, M. López-Pozo, S. K. Polutchko, W. W. Adams, Zeaxanthin, a Molecule for Photoprotection in Many Different Environments. Molecules 25, 5825 (2020).

11. C. Kulheim, J. Agren, S. Jansson, Rapid Regulation of Light Harvesting and Plant Fitness in the Field. Science (1979) 297, 91–94 (2002).

12. S. P. Long, et al., Into the Shadows and Back into Sunlight: Photosynthesis in Fluctuating Light. Annual Review of Plant Biology 73, 617–648 (2022).

13. L. Dall’Osto, S. Caffarri, R. Bassi, A mechanism of nonphotochemical energy dissipation, independent from PsbS, revealed by a conformational change in the antenna protein CP26. Plant Cell 17, 1217–32 (2005).

14. J. Kromdijk, et al., Improving photosynthesis and crop productivity by accelerating recovery from photoprotection. Science 354, 857–861 (2016).

15. de Souza AP, et al., Soybean photosynthesis and crop yield is improved by accelerating recovery from photoprotection. Science (1979) (2022).

16. R. Goss, B. Lepetit, Biodiversity of NPQ. J Plant Physiol 172C, 13–32 (2015).

17. R. Goss, D. Latowski, Lipid Dependence of Xanthophyll Cycling in Higher Plants and Algae. Front Plant Sci 11, 455 (2020).

18. G. Perin, A. Bellan, A. Bernardi, F. Bezzo, T. Morosinotto, The potential of quantitative models to improve microalgae photosynthetic efficiency. Physiologia Plantarum 166 (2019).

19. E. Sforza, D. Simionato, G. M. Giacometti, A. Bertucco, T. Morosinotto, Adjusted light and dark cycles can optimize photosynthetic efficiency in algae growing in photobioreactors. PLoS One 7, e38975 (2012).

20. L. Kalituho, J. Rech, P. Jahns, The roles of specific xanthophylls in light utilization. Planta 225, 423–39 (2007).

21. H. Hartel, H. Lokstein, B. Grimm, B. Rank, Kinetic Studies on the Xanthophyll Cycle in Barley Leaves (Influence of Antenna Size and Relations to Nonphotochemical Chlorophyll Fluorescence Quenching). Plant Physiol 110, 471–482 (1996).

22. L. Dall’Osto, et al., Two mechanisms for dissipation of excess light in monomeric and trimeric light-harvesting complexes. Nat Plants 3, 17033 (2017).

23. A. H. Short, et al., Xanthophyll-cycle based model of the rapid photoprotection of Nannochloropsis in response to regular and irregular light/dark sequences. The Journal of Chemical Physics 156, 205102 (2022).

24. M. I. S. Naduthodi, et al., CRISPR-Cas ribonucleoprotein mediated homology-directed repair for efficient targeted genome editing in microalgae Nannochloropsis oceanica IMET1. Biotechnology for Biofuels 12, 1–11 (2019).

25. Q. Wang, et al., Genome engineering of Nannochloropsis with hundred-kilobase fragment deletions by Cas9 cleavages. The Plant Journal 106, 1148–1162 (2021).

26. A. Bellan, F. Bucci, G. Perin, A. Alboresi, T. Morosinotto, Photosynthesis regulation in response to fluctuating light in the secondary endosymbiont alga nannochloropsis gaditana. Plant and Cell Physiology 61 (2020).

27. B. Bailleul, et al., An atypical member of the light-harvesting complex stress-related protein family modulates diatom responses to light. Proc Natl Acad Sci U S A 107, 18214–9 (2010).

28. G. Perin, et al., Cultivation in industrially relevant conditions has a strong influence on biological properties and performances of Nannochloropsis gaditana genetically modified strains. Algal Research 28, 88–99 (2017).

29. A. Meneghesso, et al., Photoacclimation of photosynthesis in the Eustigmatophycean Nannochloropsis gaditana. Photosynthesis Research 129, 291–305 (2016).

30. S. Park, et al., Chlorophyll – carotenoid excitation energy transfer and charge transfer in Nannochloropsis oceanica for the regulation of photosynthesis. Proc Natl Acad Sci U S A 116, 1–6 (2019).

31. S. Basso, et al., Characterization of the photosynthetic apparatus of the Eustigmatophycean Nannochloropsis gaditana: evidence of convergent evolution in the supramolecular organization of photosystem I. Biochim Biophys Acta 1837, 306–14 (2014).

32. J. S. Brown, Functional Organization of Chlorophyll a and Carotenoids in the Alga, Nannochloropsis salina. Plant Physiology 83, 434 (1987).

33. L. M. Lubián, et al., Nannochloropsis (Eustigmatophyceae) as source of commercially valuable pigments. Journal of Applied Phycology 12, 249–255 (2000).

34. E. Kress, P. Jahns, The dynamics of energy dissipation and xanthophyll conversion in arabidopsis indicate an indirect photoprotective role of zeaxanthin in slowly inducible and relaxing components of non-photochemical quenching of excitation energy. Frontiers in Plant Science 8, 2094 (2017).

35. K. K. Niyogi, O. Bjorkman, A. R. Grossman, Chlamydomonas Xanthophyll Cycle Mutants Identified by Video Imaging of Chlorophyll Fluorescence Quenching. Plant Cell 9, 1369–1380 (1997).

36. J. M. Buck, P. G. Kroth, B. Lepetit, Identification of sequence motifs in Lhcx proteins that confer qE-based photoprotection in the diatom Phaeodactylum tricornutum. The Plant Journal 108, 1721–1734 (2021).

37. T. Lacour, M. Babin, J. Lavaud, Diversity in Xanthophyll Cycle Pigments Content and Related Nonphotochemical Quenching (NPQ) Among Microalgae: Implications for Growth Strategy and Ecology. Journal of Phycology 56, 245–263 (2020).

38. L. Taddei, et al., Dynamic Changes between Two LHCX-Related Energy Quenching Sites Control Diatom Photoacclimation. Plant Physiol 177, 953–965 (2018).

39. B. Lepetit, et al., The diatom Phaeodactylum tricornutum adjusts nonphotochemical fluorescence quenching capacity in response to dynamic light via fine-tuned Lhcx and xanthophyll cycle pigment synthesis. New Phytologist 214, 205–218 (2017).

40. T. Morosinotto, R. Baronio, R. Bassi, Dynamics of chromophore binding to Lhc proteins in vivo and in vitro during operation of the xanthophyll cycle. Journal of Biological Chemistry 277, 36913–36920 (2002).

41. A. Alboresi, et al., Conservation of core complex subunits shaped the structure and function of photosystem I in the secondary endosymbiont alga Nannochloropsis gaditana. New Phytol 213, 714–726 (2017).

42. M. Havaux, K. K. Niyogi, The violaxanthin cycle protects plants from photooxidative damage by more than one mechanism. Proceedings of the National Academy of Sciences 96, 8762–8767 (1999).

43. S. Cazzaniga, et al., Domestication of the green alga Chlorella sorokiniana: reduction of antenna size improves light-use efficiency in a photobioreactor. Biotechnol Biofuels 7, 157 (2014).

44. T. De Mooij, et al., Antenna size reduction as a strategy to increase biomass productivity: a great potential not yet realized. Journal of Applied Phycology (2014) https://doi.org/10.1007/s10811-014-0427-y.

45. G. Perin, F. Gambaro, T. Morosinotto, Knowledge of Regulation of Photosynthesis in Outdoor Microalgae Cultures Is Essential for the Optimization of Biomass Productivity. Front Plant Sci 13 (2022).

46. G. Perin, A. Bernardi, A. Bellan, F. Bezzo, T. Morosinotto, A mathematical model to guide genetic engineering of photosynthetic metabolism. Metabolic Engineering 44, 337–347 (2017).

47. A. Garcia-Molina, D. Leister, Accelerated relaxation of photoprotection impairs biomass accumulation in Arabidopsis. Nature Plants 6, 9–12 (2020).

48. G. G. Lehretz, A. Schneider, D. Leister, U. Sonnewald, High non-photochemical quenching of VPZ transgenic potato plants limits CO 2 assimilation under high light conditions and reduces tuber yield under fluctuating light. Journal of Integrative Plant Biology (2022) https://doi.org/10.1111/JIPB.13320 (July 7, 2022).

49. G. Perin, et al., Generation of random mutants to improve light-use efficiency of Nannochloropsis gaditana cultures for biofuel production. Biotechnol Biofuels 8, 161 (2015).

50. A. R. Wellburn, The spectral determination of chlorophylls a and b, as well as total carotenoids, using various solvents with spectrophotometers of different resolution. Journal of Plant Physiology 144, 307–313 (1994).

51. A. Färber, P. Jahns, The xanthophyll cycle of higher plants: influence of antenna size and membrane organization. Biochimica et Biophysica Acta (BBA) - Bioenergetics 1363, 47–58 (1998).

52. S. W. Jeffrey, R. F. C. Mantoura, S. W. Wright, Phytoplankton pigments in oceanography: guidelines to modern methods. Monographs on Oceanographic Methodology (1997).

53. K. Maxwell, G. N. Johnson, Chlorophyll fluorescence - A practical guide. Journal of Experimental Botany 51, 659–668 (2000).

54. N. R. Baker, Chlorophyll fluorescence: a probe of photosynthesis in vivo. Annu Rev Plant Biol 59, 89–113 (2008).

