## Supplementary Information for "Modulation of xanthophyll cycle impacts biomass productivity in the marine microalga *Nannochloropsis*"

Supporting Information Text

**Supporting results**

***Impact of alterations in the xanthophyll cycle on photosynthetic functionality***

The Chl fluorescence kinetics after exposure to increasing light intensities was also monitored for all strains to assess in more detail the impact of alterations in the xanthophyll cycle on the functionality of the photosynthetic machinery. This confirmed a small reduction in NPQ of the ZEP over-expressor with respect to WT, while *vde KO* and *lhcx1 KO* showed a total absence of response (Supplementary figure S5a). In all strains, the photosynthetic electron transport (ETR) increased as a function of light intensity, reaching saturation at approx. 500 μmol photons m^−2^ s^−1^. When light intensity further increased, ETR decreased suggesting cells cannot process efficiently all the received photons (Supplementary figure S5b). In *vde KO* ETR values remained lower than the parental strain. *lhcx1 KO* strain showed instead an ETR higher than WT at saturating light intensities. The greatest increase in ETR was observed for the *ZEP* OE, that also reached saturation at higher light intensities than the parental strain (Supplementary figure S5b).

The fluorescence parameter qL can also be exploited to assess photochemical activity: when all reaction centers are available for photochemical reactions its value is 1, whilst it trends to 0 when the photochemical capacity is saturated (1). Both *vde KO* and *lhcx1 KO* showed a faster reduction of qL as the light intensity increases, suggesting their reactions centers were more easily saturated. The *ZEP* OE, instead, showed a higher photochemical activity than the parental strains at saturating light intensities (Supplementary figure S5c). This is also confirmed by the trend of PSII quantum yield of samples illuminated with increasing irradiances, where the *ZEP* OE showed a slower reduction of PSII activity with respect to the parental strain (Supplementary figure S5d).

The photosynthetic electron transport activity was assessed also using an alternative method, measuring the oxygen evolution upon exposure to increasing light intensity. Whilst no difference between the *ZEP* OE and the parental strain was observed, both *vde KO* and *lhcx1 KO* showed instead a strong reduction (Supplementary figure S5e), highlighting the importance of NPQ and xanthophyll cycle to preserve photosynthetic functionality in cells exposed to over-saturating irradiances.

**Supporting materials and methods**

***Design of the vector for the overexpression of genes of interest in Nannochloropsis***

A modular vector to enable effective expression of genes of interest (GOI) in *Nannochloropsis* was developed, fusing a cassette conferring resistance to Zeocin (2), including the *Sh-ble* gene under the control of the endogenous constitutive UBIQUITIN promoter and the FCPA terminator from *P. tricornutum* (3), with a second cassette enabling the expression of specific GOIs. To achieve the desired vector modularity, all the regulatory elements controlling the expression cassette were surrounded by unique restriction sites, in order to facilitate the replacement of GOIs (Supplementary figure S10a). The endogenous LIPID DROPLET SURFACE PROTEIN promoter (*LDSP*, Gene ID: 100086g4) was chosen as a strong regulatory element to drive the expression of the GOI, according to the expression of the *LDSP* gene that shows a four-fold higher value with respect to the endogenous constitutive UBIQUITIN promoter (4) (Supplementary figure S10b), together with the FCPA terminator from *P. tricornutum*. Although the functionality of the *LDSP* promoter to drive the expression of GOIs was already assessed in *Nannochloropsis oceanica* (5)*,* its activity in *N. gaditana* has not been investigated yet. The region 5’-upstream of the start codon of the *LDSP* gene was chosen as the *LDSP* promoter and the region extending to the next upstream gene (Gene ID: Naga_1000086g5) was cloned. When this sequence was used to drive the expression of the cassette conferring resistance to Zeocin, no colonies were obtained. Repeated rounds of transformation with progressively longer versions of the *LDSP* promoter extending towards the 3’, up to include the first exon and intron of the *LDSP* gene, led to the identification of the sequence to get full functionality. PlantCARE (6) indeed predicted three TATA boxes in the final functional version of the promoter, belonging to the first intronic sequence, which might explain why the shorter versions of the promoter are not functional. The luciferase gene from *Renilla reniformis* and codon optimized for *Chlamydomonas reinhardtii* (7), *Crluc*, was used as a reporter gene to test the functionality of the expression cassette (Supplementary figure S10c). Zeo^R^ colonies were obtained and the presence of the *Crluc* gene integrated in the DNA was confirmed via colony PCR (Supplementary figure S10d). Luciferase activity was confirmed by monitoring the luminescence signal, validating the functionality of the expression cassette designed in this work in *Nannochloropsis* (Supplementary figure S10e).

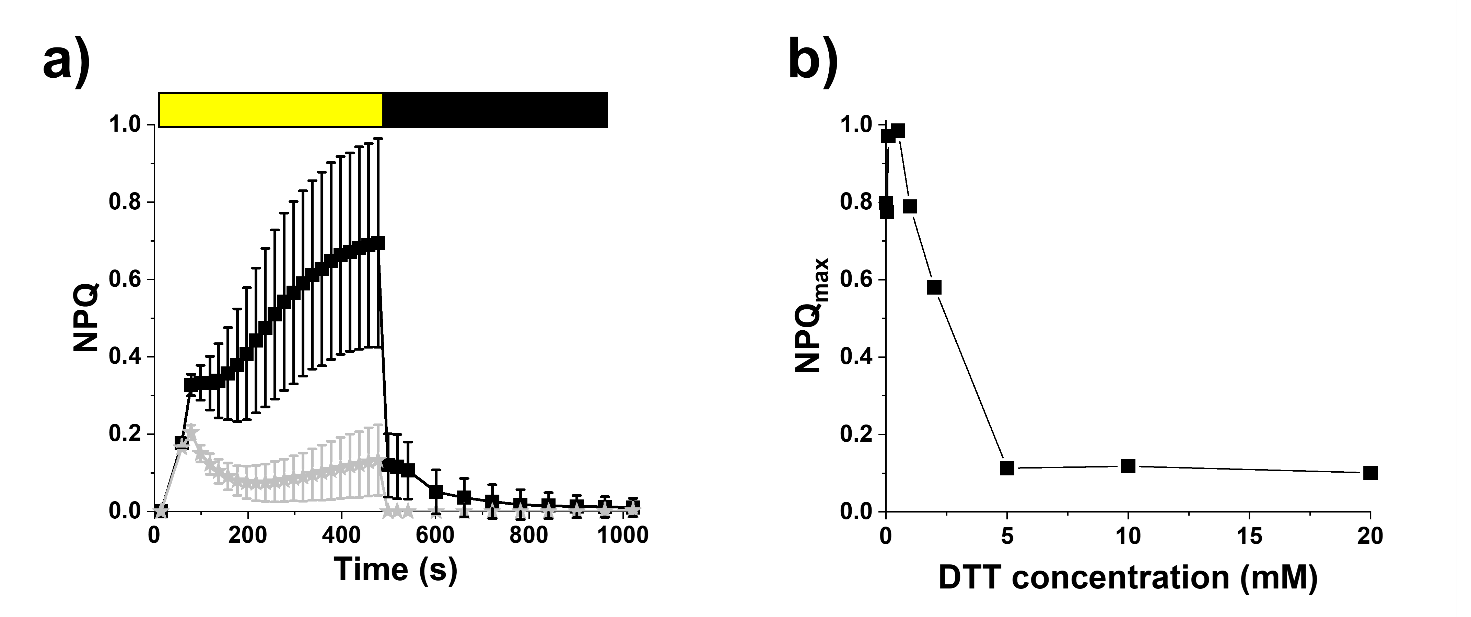

Fig. S1. Influence of zeaxanthin on NPQ activation in *Nannochloropsis*. a) NPQ induction analysis of *Nannochloropsis* cells grown at 100 μmol photons m^−2^ s^−1^. Induction kinetics of untreated (black squares) and 20 mM DTT-treated (gray stars) samples are shown. The induction protocol consists of 8 min light at 800 μmol photons m^−2^ s^−1^ (yellow box) and 15 min of dark recovery (black box). Data are reported as average ± SD of 5 biological replicates. b) Titration of DTT impact on NPQ. NPQ is here expressed as maximal activity as a function of the concentration of DTT.

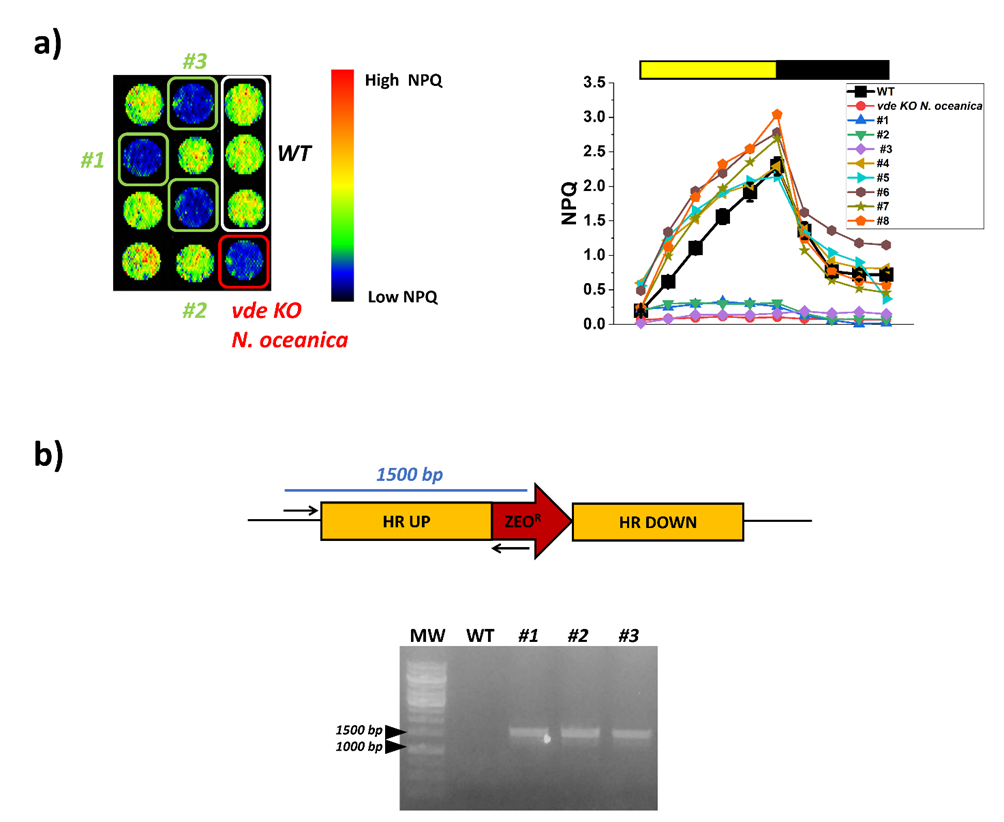

Fig. S2. Isolation of *vde KO* mutant in *Nannochloropsis gaditana*. VDE catalyses the conversion of violaxanthin to zeaxanthin, triggering the activation of Non-Photochemical Quenching (NPQ). a) KO mutants for the VDE protein (GENE ID: Naga100041g46) show a defect in the NPQ response, as observed for the *vde KO* mutant of *N. oceanica*, used as control (8). On the left of panel a), a chlorophyll fluorescence image of an agar plate containing 12 spots corresponding to 8 mutant strains of *N. gaditana*, three copies of its parental strain (WT) and the *vde KO* mutant strain of *N. oceanica* is presented. *N. gaditana* strains #1, #2 and #3 show the photosynthetic phenotype of potential *vde KO* strains, as indicated by the lack of the NPQ response. Yellow and black boxes in the right panel a) indicate saturating light and dark exposure to investigate NPQ activation and relaxation kinetics, respectively.

b) *N. gaditana* strains #1, #2 and #3 resulted in genuine *vde KO* strains after validation of their genotype via colony PCR, by amplification of the left border of the integration locus. Primers For: CTGCTCCTCCCATTTCCCATG and Rev: GCATAATTAAAGCTATTCGGTCCAATTG used in the colony PCR of panel b anneal on the genomic sequence upstream of the 5’-homology region and on the Zeocin resistance cassette, respectively. Amplification is expected only if cassette is inserted in the expected genomic region. WT, wild-type.

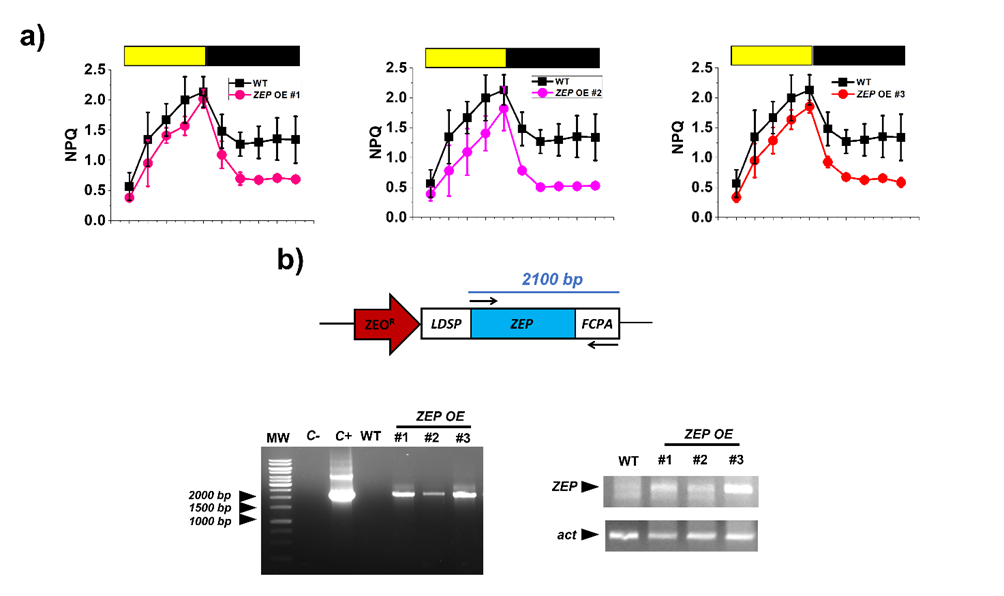

**Fig. S3. Isolation of *ZEP OE* strain in *Nannochloropsis gaditana*.** ZEP catalyses the conversion of zeaxanthin to violaxanthin, enabling the relaxation of Non-Photochemical Quenching (NPQ). A mutant strain overexpressing ZEP (*ZEP OE*) should show a faster relaxation of NPQ than the parental strain. a) NPQ kinetic of potential *ZEP OE* strains of *N. gaditana* during screening of a population of over-expressing strains, where a reduced NPQ activation and a faster relaxation is observed in mutants #1, #2 and #3 with respect to the parental strain. Yellow and black boxes indicate saturating light and dark exposure to investigate NPQ activation and relaxation kinetics, respectively. b) The genotype of the three strains was validated via colony PCR, by amplification of the portion of the cassette carrying the ZEP coding sequence together with the FCPA terminator. Primers For: ATGTTTTTCTTTTCTCAGACGT and Rev: TCAGTTGGGTTGCAACTGT used in the colony PCR. Strains #1, #2 and #3 resulted genuine *ZEP OE* strains of *N. gaditana* after validation of ZEP overexpression with respect to the parental strain via Reverse Transcriptase PCR (RT-PCR), by amplification of ZEP coding sequence from the same amount of cDNA, as indicated by the similar amplification of the housekeeping gene for ACTIN (act). Strain #3 was chosen to pursue the experiments of this work as it showed the greatest overexpression of the ZEP gene.

Primers for ZEP amplification,

For: AGGTATGGTGCAACGTCTGG and Rev: CTGGCAGTACCACTTGTTCG

and primers for act amplification,

For: ATGGCGGAAGAAGATGTGCA and Rev: GTACAGGTCCTTGCGGATGT

used in the RT-PCR. WT, wild-type; C-, negative control using water as template; C+, positive control using the plasmid carrying the ze overexpression cassette.

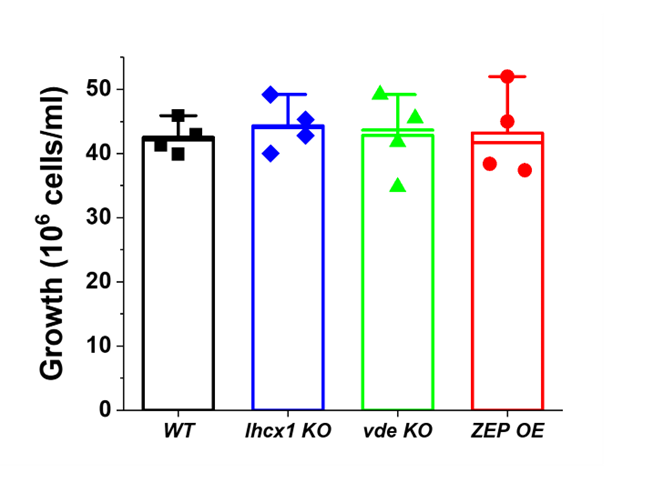

**Fig. S4. Growth of the *N. gaditana* strains investigated in this work after 4 days in Erlenmeyer ﬂasks.** Strains were cultivated in F/2, supplemented with 10 mM NaHCO_3_ to avoid carbon limitation, for 4 days, starting from a cell concentration of 5·10^6^ cells/ml (see material and methods for details).

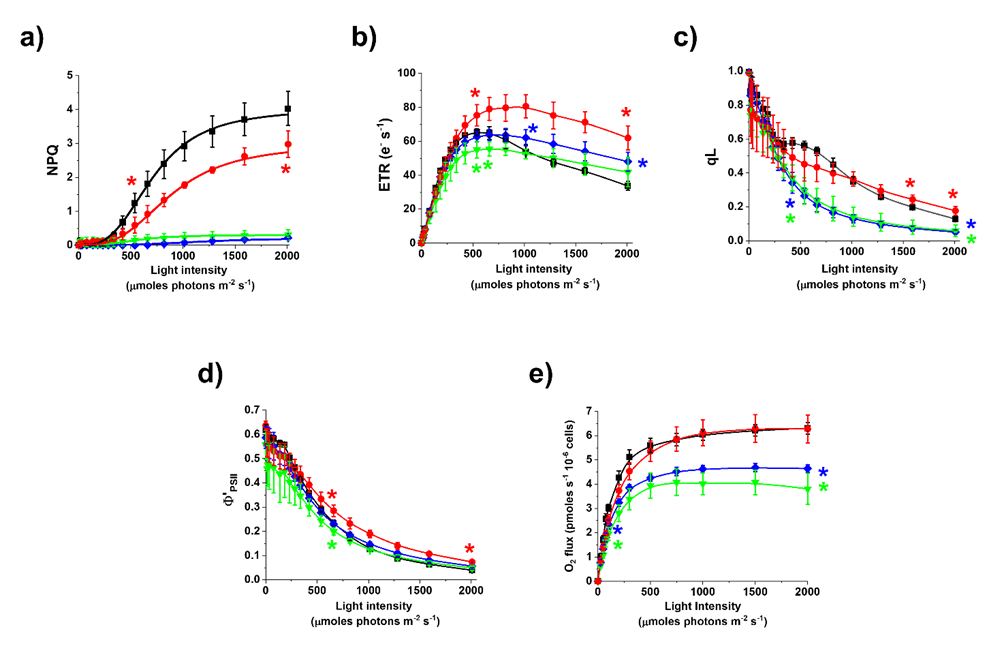

**Fig. S5. Impact of the xanthophyll cycle on photosynthesis.** The impact on photosynthetic functionality was assessed monitoring Chl fluorescence kinetics in vivo of liquid cultures cultivated in optimal light for 4 days (see materials and methods for details). a) NPQ activation, b) Photosynthetic electron transport (ETR), c) photochemical capacity (qL), d) PSII quantum yield of illuminated cells (Φ’ _PSII_) and e) oxygen evolution at increasing irradiances. The same number of cells was used for the three measurements. WT *Nannochloropsis* strain, black squares; *lhcx1 KO*, blue diamonds; *vde KO*, green downward triangles; ZEP overexpressing strain, red circles. Data are expressed as average ± SD of four independent biological replicates. Asterisks indicate statistically significant differences between mutants and parental strain (One-way ANOVA, p-value<0.05). For each strain, all the data points between the two asterisks show statistically significant difference with respect to the WT.

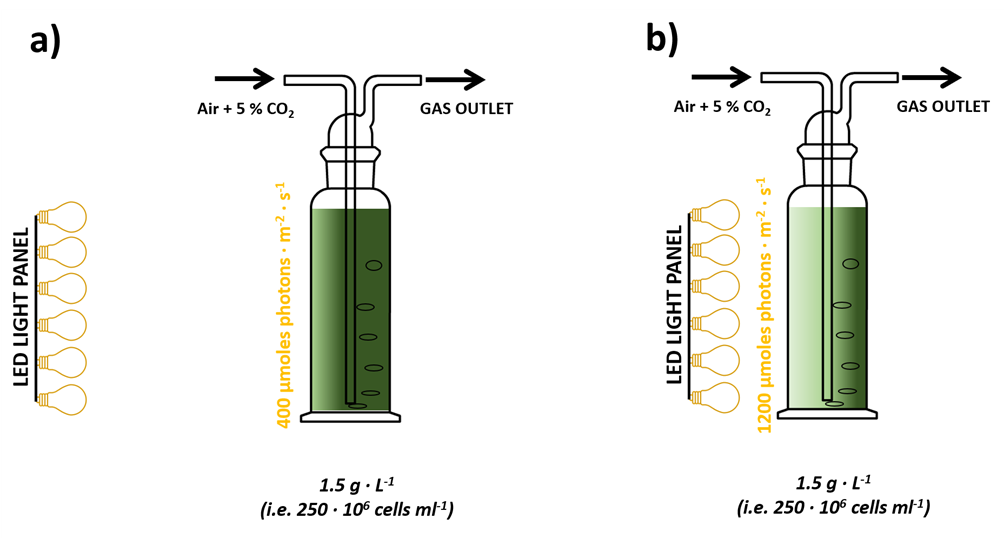

**Fig. S6. Experimental set-up for semi-continuous cultures.** *Nannochloropsis* cultivation was performed in Drechsel bottles starting from 1.5 g L^-1^ biomass concentration and cultures were diluted every other day to restore this value. Mixing and carbon source was provided through the insufflation of air enriched with 5% CO_2_ (v/v), whilst light energy was provided with a LED panel from one side of the culture to get 400 and 1200 µmol photons m^-2^ s^-1^, as indicated in a) and b), respectively.

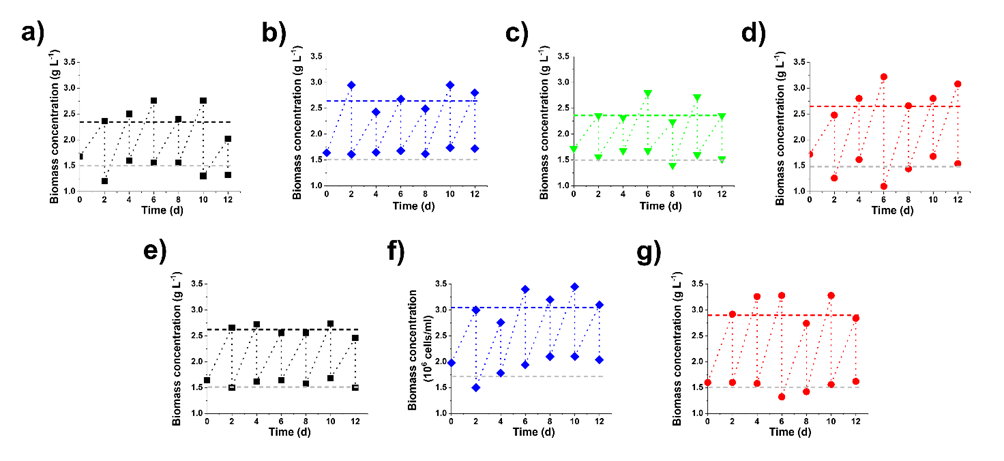

**Fig. S7. *Nannochloropsis* semicontinuous cultures.** Data were collected before and after dilution to restore the intial biomass concentration of 250 · 10^6^ cells/ml, corresponding to 1.5 g L^-1^, for *N. gaditana* WT (black squares, a and e), *lhcx1 KO* (blue diamonds, b and f), *vde KO* (green downward triangles, c) and ZEP over-expressor (red circles, d and g). Data at 400 and 1200 µmol photons · m^-2^ · s^-1^ are reported in a, b, c, d and e, f, g, respectively. Data for the *vde KO* at higher irradiance are not reported as the strain died in these conditions.

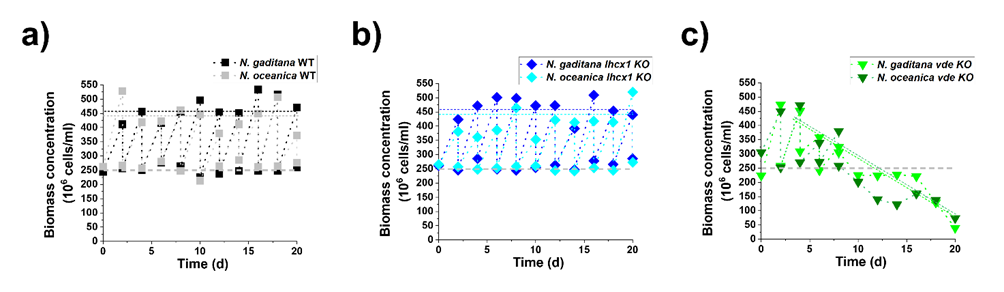

**Fig. S8. Comparison between semi-continuous cultures of *N. gaditana* and *N. oceanica* strains.** Data were collected before and after dilution to restore the intial biomass concentration of 250 · 10^6^ cells/ml, corresponding to 1.5 g L^-1^ for the two parental strains (WT, a), *lhcx1 KO* (b) and *vde KO* strains (c). These data come from semi-continuous cultures at 1200 µmol photons m^-2^ s^-1^.

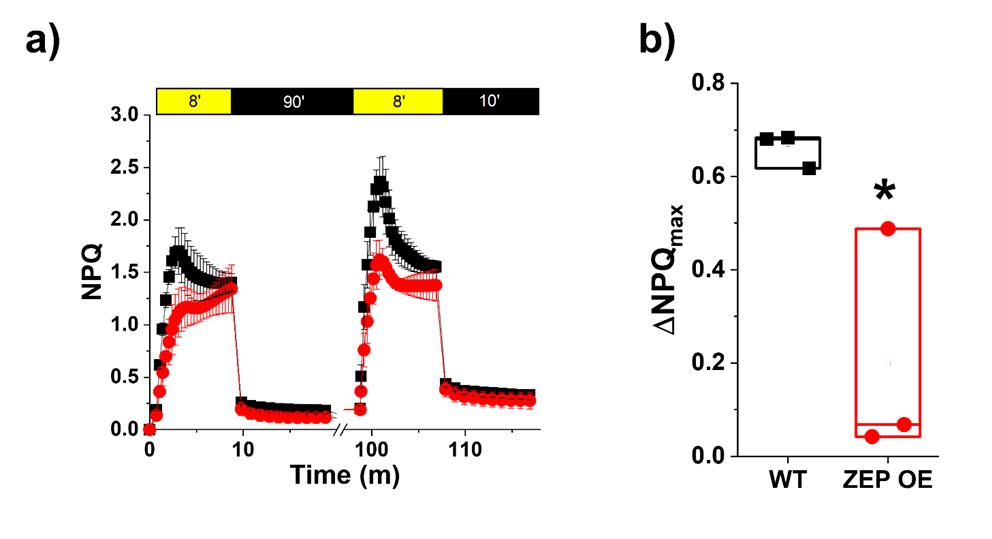

**Fig. S9. Effect of ZEP over-expression on the contribution of zeaxanthin on NPQ.** a) Chlorophyll fluorescence kinetics upon exposure of *Nannochloropsis gaditana* WT (black squares) and ZEP over-expressor (red circles) to two repetitions of 8 minutes light interspersed by 90 minutes dark. Yellow and black boxes indicate light and dark intervals, respectively. b) Difference in the maximum NPQ value between the second and first light treatment. Data are expressed as average ± SD of three independent biological replicates. Asterisk indicates statistically significant difference between ZEP over-expressor and parental strain (One-way ANOVA, p-value<0.05).

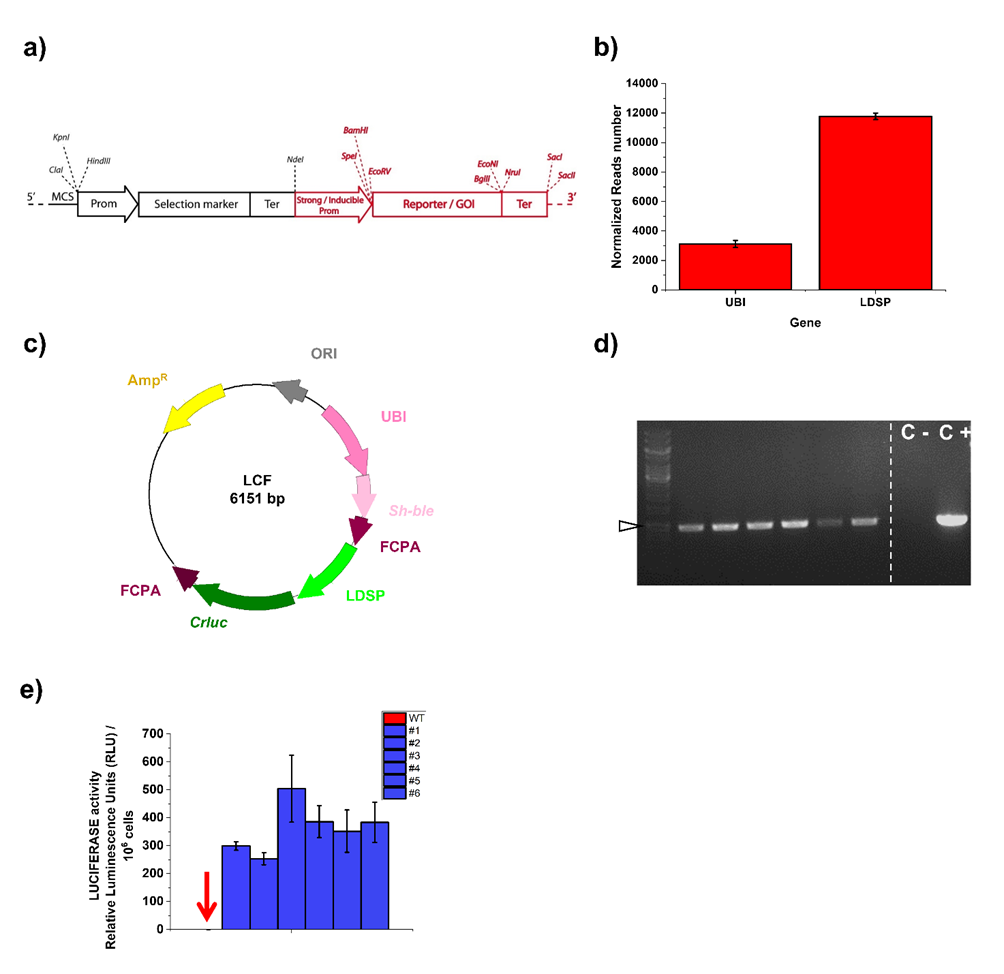

**Fig. S10. Design of the construct for the overexpression of genes of interest in *Nannochloropsis*.** a) Schematic overview of the construct, where the cassettes used for transformants selection (in black) and proteins over-expression (in red) are highlighted. Each element of the over-expression cassette was designed to be surrounded by unique restriction sites. This feature allows the simple opening of the vector for investigating molecular regulatory regions and for the replacement of the coding sequence of specific target proteins. Prom, Promoter; Ter, Terminator; GOI, Gene Of Interest. b) Normalized reads number for the UBIQUITIN (UBI) and LIPID DROPLET SURFACE PROTEIN (LDSP) genes from *N. gaditana* cells acclimated to 100 µmol photons m^-2^ s^-1^. Data come from average ± SD of 3 biological replicates. c) Schematic overview of the vector (LCF) used to study the functionality of the LDSP promoter in *N. gaditana*. The backbone comes from the pBlueScript II SK (+) vector. The molecular components are depicted in different colors: yellow, Amp^R^; grey, E. coli ORI; pink, endogenous UBIQUITIN promoter (UBI); light-pink, Sh-ble gene conferring resistance to Zeocin; violet, FCPA terminator; light-green, LDSP promoter; green, CrLUC gene. For ZEP overexpression, CrLUC was replaced with the endogenous ZEP coding sequence. d) *N. gaditana* transformants were tested via colony PCR, to validate the successful integration of the CrLUC gene (936 bp) in the genome. Six independent transformants (lanes 2 – 7) confirmed the presence of the CrLUC gene in their genome with respect to WT strains (C -). C -, WT *N. gaditana* strain; C +, plasmid DNA (LCF vector). The left and the right part of the picture come from different regions of the same agarose gel. The first lane indicates the molecular weight (MW) and the arrow indicates 1000 bp. e) LUCIFERASE activity of the six strains analysed in d). LUCIFERASE activity is expressed as Relative luminescence units (RLU) per million of cells. *N. gaditana* WT is indicated on the left of the series by a red arrow and measured values are negligible. Data are expressed as averages and SD of 3 biological replicates and indicate the functionality of the designed vector for effective protein overexpression in *N. gaditana*.

Table S1. Pigment content of *Nannochloropsis gaditana* after 2 h exposure to limiting (LL) and excess (EL) light conditions. Carotenoids are reported as mol/100 mol of Chl or, in the last three columns, normalized over the sum of the three xanthophylls violaxanthin (Vx), antheraxanthin (Ax) and zeaxanthin (Zx)(VAZ). Data are expressed as the average ± SD of 3 independent biological replicates. Asterisks indicate statistically significant differences between EL and LL conditions (One-way ANOVA, p-value<0.05). Vaucheriax, Vaucheriaxanthin.

|  | Vx | Vaucheriax | Ax | Zx | β–car | % Vx/VAZ | % Ax/VAZ | % Zx/VAZ |
| --- | --- | --- | --- | --- | --- | --- | --- | --- |
| LL | 24.37 ± 2.41 | 4.88 ± 0.61 | 0.37 ± 0.53 | 0.99 ± 0.15 | 1.67 ± 2.37 | 93.84 ± 1.90 | 3.30 ± 0.33 | 3.96 ± 0.28 |
| EL | 18.12 ± 0.52* | 5.09 ± 0.76 | 4.69 ± 1.37* | 3.45 ± 0.83* | 1.17 ± 1.66 | 70.32 ± 4.09* | 17.72 ± 3.32* | 13.29 ± 1.47* |

Table S2. Pigment content of *Nannochloropsis gaditana* strains investigated in this work. Data here reported come from cultures after 4 days of growth in optimal light (OL), 2 h treatment with excess light (HL) and 1.5 h recovery in optimal light (ROL). Carotenoids are reported as mol/100 mol of Chl. Data are expressed as the average ± SD of 4 independent biological replicates. Vaucheriax, Vaucheriaxanthin.

|  | Vaucheriax | | | β–car | | |
| --- | --- | --- | --- | --- | --- | --- |
|  | **OL** | **HL** | **ROL** | **OL** | **HL** | **ROL** |
| WT | 21 ± 1.8 | 21 ± 1.85 | 21.5 ± 2.2 | 0.51 ± 0.37 | 0.48 ± 0.23 | 0.6 ± 0.7 |
| *lhcx1 ko* | 20 ± 2 | 20.4 ± 2.5 | 20.6 ± 3 | 0.75 ± 0.41 | 1.23 ± 0.76 | 1.2 ± 0.7 |
| *vde KO* | 21.2 ± 2.6 | 20.6 ± 1.8 | 24.5 ± 1.6 | 0.69 ± 0.5 | 0.58 ± 0.32 | 0.8 ± 0.6 |
| *ZEP OE* | 20 ± 1.5 | 19.5 ± 1.8 | 20 ± 2.8 | 0.8 ± 0.66 | 0.75 ± 0.35 | 1.26 ± 1 |

**Table S3. Pigment content (Chl concentration in picograms per cell and Chl/Car) of the strains investigated in this work.** Low and High irradiance correspond to 400 and 1200 µmol photons · m^-2^ · s^-1^, respectively, n > 10.

| Species | Strain | Low Irradiance | | High Irradiance | |
| --- | --- | --- | --- | --- | --- |
|  |  | **Chl (pg/cell)** | **Chl/Car** | **Chl (pg/cell)** | **Chl/Car** |
| *N. gaditana* | **WT** | 0.14 ± 0.01 | 2.73 ± 0.09 | 0.07 ± 0.004 | 2.29 ± 0.16 |
| *N. gaditana* | ***lhcx1 KO*** | 0.13 ± 0.02 | 3.12 ± 0.21 | 0.05 ± 0.004 | 2.21 ± 0.03 |
| *N. gaditana* | ***vde KO*** | 0.11 ± 0.01 | 2.9 ± 0.1 | / | / |
| *N. gaditana* | ***ZEP OE*** | 0.16 ± 0.02 | 2.74 ± 0.04 | 0.05 ± 0.004 | 2.17 ± 0.11 |
| *N. oceanica* | **WT** | 0.13 ± 0.02 | 2.93 ± 0.24 | 0.05 ± 0.008 | 2.05 ± 0.11 |
| *N. oceanica* | ***lhcx1 KO*** | 0.1 ± 0.01 | 3.08 ± 0.22 | 0.05 ± 0.003 | 2.2 ± 0.13 |
| *N. oceanica* | ***vde KO*** | 0.13 ± 0.03 | 2.91 ± 0.08 | / | / |

**Table S4. Maximal photosynthetic efficiency (Φ_PSII_) of the strains investigated in this work.** Low and High irradiance correspond to 400 and 1200 µmol photons · m^-2^ · s^-1^, respectively, n > 10.

| Species | Strain | Low Irradiance | High Irradiance |
| --- | --- | --- | --- |
| *N. gaditana* | **WT** | 0.63 ± 0.04 | 0.47 ± 0.04 |
| *N. gaditana* | ***lhcx1 KO*** | 0.64 ± 0.02 | 0.5 ± 0.02 |
| *N. gaditana* | ***vde KO*** | 0.6 ± 0.01 | / |
| *N. gaditana* | ***ZEP OE*** | 0.62 ± 0.01 | 0.4 ± 0.04 |
| *N. oceanica* | **WT** | 0.67 ± 0.04 | 0.48 ± 0.06 |
| *N. oceanica* | ***lhcx1 KO*** | 0.69 ± 0.03 | 0.46 ± 0.03 |
| *N. oceanica* | ***vde KO*** | 0.69 ± 0.02 | / |

**Table S5. Photosynthetic activity expressed as oxygen evolution rate during the first fluctuation cycle of the protocol of figure 6a.** The oxygen evolution rate is expressed as pmol O_2_ s^-1^ 10^-6^ cells for all strains investigated in this work at 100, 300 and 15 µmol photons · m^-2^ · s^-1^. Asterisks indicate statistically significant differences between mutants and parental strain (One-way ANOVA, p-value<0.05).

| Strain | Irradiance | | |
| --- | --- | --- | --- |
|  | **100** | **300** | **15** |
| WT | 9.3 ± 1.4 | 12.45 ± 4.5 | 1.31 ± 0.17 |
| *lhcx1 KO* | 5.74 ± 1.63* | 7.26 ± 2.7* | 0.63 ± 0.19* |
| *vde KO* | 12.7 ± 1.75 | 16.3 ± 7.45 | 1.35 ± 0.45 |
| *ZEP OE* | 9.5 ± 2.5 | 12.9 ± 5.3 | 1.19 ± 0.28 |

**Table S6. Mathematical description of the oxygen evolution trends over the cycles of light fluctuation described in figure 6, for the *N. gaditana* strains investigated in this work.** Trends observed in this experiment follow a linear function (y = b + ax) and the corresponding parameters for the different strains of this work are here reported. Data are expressed as the average ± SD of 4 independent biological replicates. Asterisks indicate when the slope of the linear function is statistically significant different from zero (One-way ANOVA, p-value < 0.05).

|  | 300 | | | | 15 | | | |
| --- | --- | --- | --- | --- | --- | --- | --- | --- |
|  | **b** | **a** | **Pearson’s R** | **R-Square** | **b** | **a** | **Pearson’s R** | **R-Square** |
| WT | 15.07 ± 0.35 | -0.59 ± 0.06* | -0.97 | 0.94 | 1.82 ± 0.04 | -0.5 ± 0.01* | -0.99 | 0.99 |
| *lhcx1 KO* | 7.8 ± 0.41 | 0.14 ± 0.08 | 0.56 | 0.32 | 0.85 ± 0.07 | -0.11 ± 0.02* | -0.94 | 0.88 |
| *vde KO* | 16.25 ± 0.5 | -0.03 ± 0.07 | -0.18 | 0.03 | 2 ± 0.2 | -0.48 ± 0.05* | -0.97 | 0.94 |
| *ZEP OE* | 14.8 ± 0.65 | -0.3 ± 0.12 | -0.7 | 0.48 | 1.83 ± 0.21 | -0.35 ± 0.05* | -0.93 | 0.87 |

**SI References**

1.	N. R. Baker, Chlorophyll fluorescence: a probe of photosynthesis in vivo. *Annu Rev Plant Biol* **59**, 89–113 (2008).

2.	G. Perin, *et al.*, Generation of random mutants to improve light-use efficiency of Nannochloropsis gaditana cultures for biofuel production. *Biotechnol Biofuels* **8**, 161 (2015).

3.	R. Radakovits, *et al.*, Draft genome sequence and genetic transformation of the oleaginous alga Nannochloropis gaditana. *Nat Commun* **3**, 686 (2012).

4.	A. Alboresi, *et al.*, Light Remodels Lipid Biosynthesis in Nannochloropsis gaditana by Modulating Carbon Partitioning between Organelles. *Plant Physiol* **171**, 2468–82 (2016).

5.	Y. Kaye, *et al.*, Metabolic engineering toward enhanced LC-PUFA biosynthesis in Nannochloropsis oceanica: Overexpression of endogenous Δ12 desaturase driven by stress-inducible promoter leads to enhanced deposition of polyunsaturated fatty acids in TAG. *Algal Research* **11**, 387–398 (2015).

6.	S. Rombauts, P. Déhais, M. Van Montagu, P. Rouzé, PlantCARE, a plant cis-acting regulatory element database. *Nucleic Acids Res* **27**, 295–6 (1999).

7.	M. Fuhrmann, *et al.*, Monitoring dynamic expression of nuclear genes in Chlamydomonas reinhardtii by using a synthetic luciferase reporter gene. *Plant Mol Biol* **55**, 869–81 (2004).

8.	S. Park, *et al.*, Chlorophyll – carotenoid excitation energy transfer and charge transfer in Nannochloropsis oceanica for the regulation of photosynthesis. *Proc Natl Acad Sci U S A* **116**, 1–6 (2019).

1. ^§^ Equal contribution
